# Stable Individual Differences Dominate Adult Brain Volume Variation Until Later Life

**DOI:** 10.1101/2025.05.26.655710

**Authors:** Edvard O. S. Grødem, Didac Vidal-Pineiro, Øystein Sørensen, David Bartrés-Faz, Andreas M. Brandmaier, Gabriele Cattaneo, Pablo F. Garrido, Richard N. Henson, Simone Kühn, Ulman Lindenberger, Bradley J. MacIntosh, Lars Nyberg, Alvaro Pascual-Leone, Stephen M. Smith, Cristina Solé-Padullés, Javier Solana-Sánchez, Leiv Otto Watne, Kristine B. Walhovd, Atle Bjørnerud, Anders M. Fjell

**Affiliations:** Center for Lifespan Changes in Brain and Cognition (LCBC), Department of Psychology, University of Oslo, Oslo, Norway; Computational Radiology and Artificial Intelligence (CRAI), Department of Radiology and Nuclear Medicine, Oslo University Hospital, Oslo, Norway; Department of Medicine, Faculty of Medicine and Health Sciences, Institute of Neurosciences, University of Barcelona, Barcelona, Spain; Institut Guttmann, Institut Universitari de Neurorehabilitació adscrit a la UAB, Badalona, Spain; The August Pi i Sunyer Biomedical Research Institute (IDIBAPS), Hospital Clinic of Barcelona, Barcelona, Spain; Center for Lifespan Psychology, Max Planck Institute for Human Development, Berlin, Germany; Department of Psychology, MSB Medical School Berlin, Berlin, Germany; Max Planck UCL Centre for Computational Psychiatry and Ageing Research, Berlin, Germany, and London, UK; MRC Cognition and Brain Sciences Unit, Department of Psychiatry, University of Cambridge, Cambridge, United Kingdom; Center for Environmental Neuroscience, Max Planck Institute for Human Development, Berlin, Germany; Department of Psychiatry and Psychotherapy, University Medical Center Hamburg-Eppendorf, Hamburg, Germany; Umeå Center for Functional Brain Imaging, Umeå University, Umeå, Sweden; Department of Medical and Translational Biology, Umeå University, Umeå, Sweden; Department of Diagnostics and Intervention, Umeå University, Umeå, Sweden; Oslo Delirium Research Group, Institute of Clinical Medicine, Campus Ahus, University of Oslo, Oslo, Norway; Department of Geriatric Medicine, Akershus University Hospital, Oslo, Norway; Hinda and Arthur Marcus Institute for Aging Research, Deanna and Sidney Wolk Center for Memory Health, Hebrew SeniorLife, Boston MA, USA; Department of Neurology, Harvard Medical School, Boston MA, USA; Fundació Institut d’Investigació en Ciències de la Salut Germans Trias i Pujol, Badalona, Spain; Oxford Centre for Functional MRI of the Brain (FMRIB/WIN), Oxford University, Oxford, United Kingdom

**Keywords:** stochastic dynamical system, brain aging, magnetic resonance imaging, methods, MRI, brain age, BAG

## Abstract

Individual differences in the volumes of brain structures are often linked to various conditions, including Alzheimer’s disease, schizophrenia, and overall brain health. However, it remains unclear to what extent these differences reflect individual levels present from young adulthood or diverging aging trajectories from later ages. In this study, we analyze the aging dynamics of the volumes of six brain structures based on magnetic resonance imaging (MRI) scans from a large cross-cohort longitudinal sample of cognitively healthy adults (n = 8,311 with 18,520 MRIs, ages from 18 to 97 years). From general assumptions about structural brain dynamics and measurement noise, a stochastic dynamical model was fitted to the data to estimate both the variability and persistence of structural changes across adulthood. Using this model, we calculated how much of the variance of volumetric differences between individuals can be attributed to stable levels from young adulthood versus systematic changes at older ages, as well as the theoretical sensitivity of longitudinal studies to detect individual differences in change. The findings were as follows: 1) Before age 60 years, inter-individual differences in neuroanatomical volumes almost exclusively reflect stable differences between individuals, while the influence from systematic differences in rate-of-change increases thereafter; up to 50 % of the variation being due to differences in change at 80 years. In contrast, ventricular volume reflects differences in change from early adulthood. 2) Current brain-age models are unlikely to be sensitive to detect differences in aging trajectories. 3) Imaging studies have low reliability in detecting inter-individual brain changes before age 60. After 60 years, the study reliability increases sharply with longer intervals between scans and more modestly with additional intermediate observations. In conclusion, our results reinforce the view that it is critical to distinguish stable early-adulthood levels from systematic differences in change when studying adult brain aging.

## 1 Introduction

Adult humans show enormous variation in the size of brain structures within the normal, healthy population. For example, a study found that young healthy adults vary in cortical gray matter volume by 52%, white matter volume by 49% and hippocampal volume by 43% compared to the mean (4 SD / mean) (Walhovd et al., 2011). These differences are very large compared to the modest amount of change over time in the same structures when measured longitudinally, which even in old age is less than 1% annually for most regions (Driscoll et al., 2009; Fjell et al., 2009; Sele et al., 2021). Thus, it is likely that a major part of volumetric differences in adulthood reflects stable, individual differences originating earlier in life (Walhovd et al., 2023). However, the continuous morphological changes throughout life also imply a balance between early adulthood levels and systematic differences in change (aging) contributions with effects of change accumulating over time potentially being more important later in life. Here, we aim to study the degree to which single observations capture early adulthood level vs. change effects in adulthood and how the relative contribution of each change throughout the lifespan.

Due to the greater availability and logistical feasibility of acquiring cross-sectional MRIs (i.e., a single measurement per person), there is considerable interest in estimating individualized aging trends based on cross-sectional observations. Although it is unclear whether these measures can capture the ongoing decline of individual brains (Raz & Lindenberger, 2011; Rohrer, 2025), such inferences are frequently drawn with respect to brain aging (Franke & Gaser, 2019), schizophrenia (Blake et al., 2023), brain health (Bashyam et al., 2020), a variety of genetic and lifestyle-related risk factors for Alzheimer’s disease (AD)(Habes et al., 2016), and even mortality (Cole et al., 2018).

The assumption that cross-sectional data can be used to assess individual differences in change can be problematic, because research has demonstrated that cross-sectional differences often do not translate into differences in change (Di Biase et al., 2023; Nyberg et al., 2010, 2021; Raz et al., 2005), with formal analysis further confirming this weak correspondence (e.g., Lindenberger et al., 2011). This issue has resurfaced with the increasing popularity of Brain Age Gap (BAG), often interpreted as a marker of accelerated brain aging and ongoing brain change. However, this interpretation has been repeatedly challenged. Studies show that baseline effects are likely to dominate the BAG signal (Smith et al., 2019, 2025), and other studies find that there are modest-to-null correlations between cross-sectional BAG and longitudinal brain change (Dörfel et al., 2023; Korbmacher et al., 2025). For example, we previously showed that having a higher BAG was not associated with a faster rate of brain aging, but rather reflected stable differences originating from very early in life (Vidal-Pineiro et al., 2021).

It is therefore critical to distinguish the contributions from the level at young adulthood vs. systematic differences in change to the total inter-individual differences in brain structure volumes. Importantly, different MRI measures capture ongoing brain changes to varying degrees, with some reflecting change clearly better than BAG (Smith et al., 2020; Vidal-Pineiro et al., 2021). Controlling for intracranial volume (ICV) eliminates a major source of stable brain volume variability (Barnes et al., 2010; Fürtjes et al., 2025; Schwarz et al., 2016), and since the relative contribution of degenerative change to cross-sectional estimates gradually increases with older age, brain volume will provide a better reflection of ongoing atrophy when controlling for ICV in older age (Schwarz et al., 2016).

The degree to which accumulation of systematic differences in change eventually manifests itself in observable cross-sectional differences depends on at least three factors (Brandmaier, Von Oertzen, et al., 2018; Hertzog, 1985):

- The magnitude of the initial differences at young adulthood (level effects).
- The magnitude of the systematic inter-individual differences in change rate (change effects).
- The time span over which the differences in change rate unfolds.

Thus, individual differences in brain structure at any age will depend on a mixture of stable differences and accumulation of systematic differences in change, and the relative contribution of each is likely to vary with age.

Estimating how variation in these changes depends on age is also important for designing imaging studies. Several studies have shown, through either simulations (Brandmaier, Von Oertzen, et al., 2018; Brandmaier et al., 2024) or empirical data (Vidal-Piñeiro et al., 2025), that detecting changes in brain volumes is difficult due to the small magnitude of the changes between sessions compared to the magnitude of the measurement noise. Because the volume change rate of brain structures probably varies with age, the expected signal from change will also vary.

Here, we propose to model the aging brain as a time-dependent stochastic dynamical system to quantify the relative contributions of levels from young adulthood vs. systematic differences in change. With this model, we analyze six major structural brain measures throughout the adult lifespan. Using stochastic dynamical models to describe systems has a long tradition in physics (Einstein, 1906; Gardiner et al., 1985), finance (Föllmer & Schied, 2011; Lee, 1992), engineering (Kalman, 1960; Welch, Bishop, et al., 1995) and neuroscience (Friston, 2010). Due to recent developments in Bayesian statistical methods (Hoffman, Gelman, et al., 2014), fitting such a system directly from data is now possible within a reasonable computational time-frame (Driver & Voelkle, 2018; Sørensen & McCormick, 2025). Importantly, the use of stochastic dynamical models explicitly captures how the change rate of each individual is changing over time, and thus sets it apart from other normative modeling approaches that only model cross-sectional data (Nobis et al., 2019) or do not model the dynamics and stochasticity of the change rate (Battaglini et al., 2019; Di Biase et al., 2023; Fujita et al., 2023; Pfefferbaum et al., 2013).

We hypothesize that the major part of structural brain differences can be explained by inter-individual differences in young adulthood, which remain stable for most of adult life, and that an increasingly larger proportion of the variance can be ascribed to individual differences in change at older ages. To our knowledge, this question has not previously been systematically investigated. We further use the dynamical model to calculate the theoretical correlation between the measurements of brain change with the true (i.e., latent) change, and from this analysis, we make recommendations for how best to design longitudinal studies to maximize sensitivity to ongoing brain change.

## 2 Methods

### 2.1 Data

We assembled data from 14 independent cohorts. In total, the combined sample comprised 8,311 cognitively healthy participants contributing 18,520 T1-weighted MRI sessions, including both crosssectional observations and longitudinal follow-up. The contributing datasets are listed in Table 1. Within each cohort, we aimed to include only observations corresponding to cognitively healthy adults, using the study-specific diagnostic frameworks and screening instruments available in that dataset. Detailed descriptions of recruitment, study protocols, and cohort-specific exclusion criteria are provided in the Supplementary Materials.

**Table 1:** List of datasets used in the study.

| Dataset | Abbreviation | Reference | N<br>subjects | N<br>sessions | M/F |
| --- | --- | --- | --- | --- | --- |
| UKBiobank | UKB | Miller et al., 2016 | 1287 | 2574 | 0.95 |
| UiO LCBC | LCBC | Walhovd et al., 2018 | 1274 | 2295 | 0.55 |
| OASIS-3 | OASIS-3 | LaMontagne et al., 2019 | 974 | 2120 | 0.75 |
| The Barcelona Brain Health Initiative | BBHI | Cattaneo et al., 2018 | 930 | 1201 | 1.05 |
| The Alzheimer's disease neuroimaging initiative | ADNI | Mueller et al., 2005 | 769 | 3316 | 0.83 |
| Cambridge Centre for Ageing Neuroscience | Cam-CAN | Shafto et al., 2014 | 638 | 905 | 0.99 |
| The Berlin Aging Study II | BASE-II | Bertram et al., 2014 | 430 | 730 | 1.62 |
| The Vietnam Era Twin Study of Aging | VETSA | Kremen et al., 2019 | 406 | 1453 | Male only |
| The Wayne State longitudinal data set | Wayne | Burgmans et al., 2010; Kennedy and Raz, 2009 | 313 | 497 | 0.49 |
| PREVENT-AD | Prevent-AD | Tremblay-Mercier et al., 2021 | 305 | 1360 | 0.43 |
| The BETULA project | BETULA | Nilsson et al., 2004; Nyberg et al., 2020 | 303 | 587 | 0.98 |
| The University of Barcelona | UB | Abellana-Pérez et al., 2019; Rajaram et al., 2017; Uribe et al., 2016; Vidal-Piñeiro et al., 2014 | 288 | 421 | 0.56 |
| Harvard Aging Brain Study* | HABS | Dagley et al., 2017 | 281 | 673 | 0.69 |
| COGNORM | COGNORM | Idland et al., 2017 | 113 | 388 | 0.88 |
| Combined sample |  |  | 8311 | 18520 | 0.89 |

The datasets span multiple countries and imaging centers, with age coverage from 18 to 97 years. The geographic distribution, as well as the age distribution and number of sessions contributed by each dataset, are summarized in Fig. 1. Because the aggregated sample contains a mixture of single-session participants and individuals with repeated scans, our analysis explicitly accommodate both cross-sectional and longitudinal sampling.

**Figure 1:**
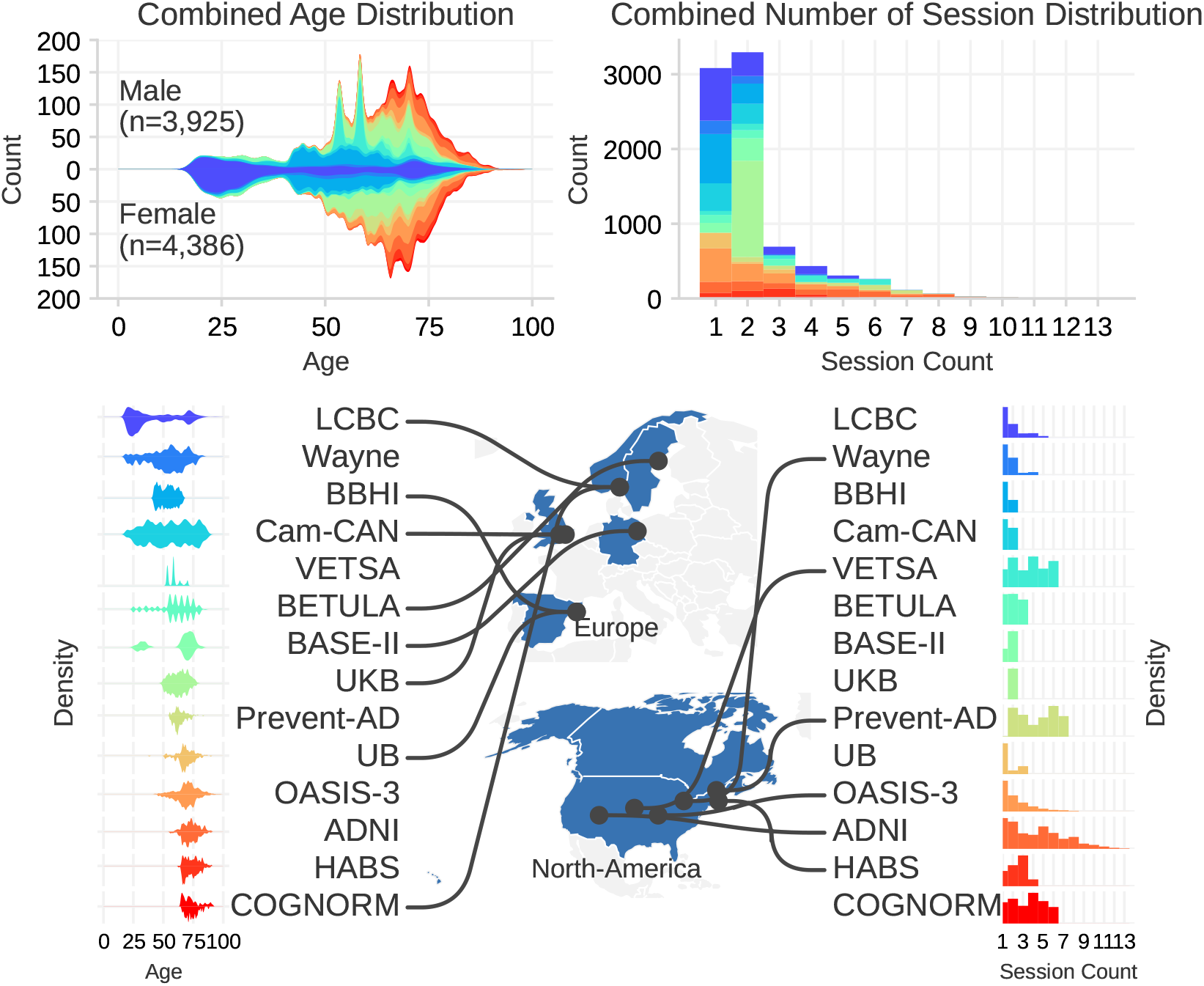
Left: Age distribution of the combined sample. The histogram shows the number of male and female participants at each age, with colors denoting different datasets. Right: The distributions of the number of sessions for each individual in the datasets.

The T1-weighted MRI scans were processed using the longitudinal stream in FreeSurfer v7.1.0 (Reuter et al., 2012). From the processed outputs, we extracted ASEG-derived (Fischl et al., 2002) volumetric measures, including the combined (left + right) hippocampal volume, total gray matter, total white matter, supratentorial volume (excluding ventricles), combined (left + right) lateral ventricles, and brainstem. To account for scanner and site-related variability and bias, the imaging site was included as a covariate in our model. In the ADNI cohort, some sites have very few participants; therefore, we constructed combined sites for ADNI based on scanner type, reducing the number of imaging sites from 125 to 39.

### 2.2 Model

#### 2.2.1 Informal Description

No study follows individuals longitudinally from early adulthood to old age. Therefore, we construct a dynamical model to understand brain development over the lifespan. Our approach is based on several assumptions that enable us to characterize the evolution of brain structure volumes under both systematic trends and random variations. Brain structures, such as the hippocampus, change gradually over time as an individual ages; abrupt changes in volume are not expected. We also expect the change rate of each structure to be stable over a short timespan such as a few years. However, over a lifetime, an individual may change their change rate relative to the population due to external factors such as lifestyle, illness, or other random influences. This is not captured by most commonly used models. To capture the changing nature of change rate, modeling the acceleration is necessary. The acceleration is assumed to have a random individualized component and a systematic influence dependent on age. An individual’s change rate at any age reflects the accumulated effect of the acceleration up to that age. The acceleration is modeled to be linearly dependent on the volume and change rate, which influence the dynamics over time. There is evidence that ICV, and by extension, brain volume, has increased over the last century (DeCarli et al., 2024; Kim et al., 2018). We adjust for ICV in our dynamical model by letting the volume at age 18 be linearly dependent on the ICV. Adjusting for ICV may also correct for other factors such as environmental factors during development and genetics that might bias some of the datasets. The measurements of brain structures are subject to considerable noise and site bias, especially compared to the subtle true changes observed over a few years (Narayanan et al., 2020; Vidal-Piñeiro et al., 2025). We therefore explicitly incorporate measurement noise and site bias into our model. Each imaging site is assumed to have an independent noise magnitude and measurement bias. The mean of the site noise is used for displaying the results. With these assumptions, the brain development of each volume is modeled as a stochastic dynamical system evolving under the influence of both systematic trends and random factors.

#### 2.2.2 Formal Description

We model the system dynamics using the latent state ***z***(*t*) defined as:

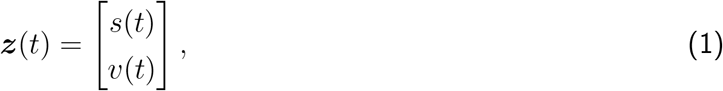

with *s*(*t*) representing the true volume of the brain structure when removing measurement noise and site bias, and *v*(*t*) representing the first derivative with respect to time of the volume, which we refer to as the change rate. The time variable *t* denotes the age of the individual, with an initial age of *t*_0_ set to the age of 18 years. Based on the assumption in the previous section, the stochastic time-dependent dynamical model is given by

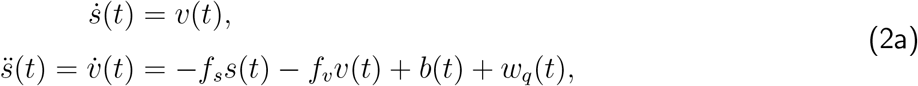

where (·) and (··) denote the first and second time derivative operation. *f*_*s*_ is the feedback coefficient from the volume and *f*_*v*_ is the feedback coefficient from the change rate. To prevent unstable exponential growth of the structures, which may not be biologically plausible, the feedback is restricted to be negative. *b*(*t*) is an age dependent bias, and *w*_*q*_ is Gaussian white noise with an age dependent variance 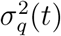.

The observed volume measurement *ŝ*(*t*) is modeled as,

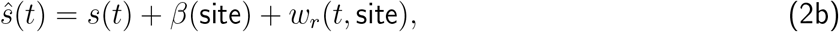

where *β*(site) is a site dependent-bias term and *w*_*r*_(*t*, site) is Gaussian white noise with site-dependent variance 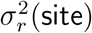. *β*(site) accounts for different imaging sites having different scanners and scanning protocols that may cause a bias in the volume measurement (X. Han et al., 2006) and 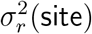 account for that some sites have more reliable scanners than others. *β* is constrained such that ∑_*j*∈sites_ *β*_*j*_ := 0, ensuring that *ŝ*(*t*) is an unbiased estimator of *s*(*t*). At time *t*_0_ the population latent volume and change rate are modeled as a multivariate Gaussian distribution,

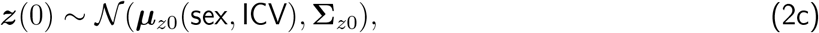

where the initial mean ***µ***_*z*0_ is assumed to be linearly dependent on both the sex and the ICV of the individuals and **Σ**_*z*0_ is the covariance matrix of the initial distribution.

In summary, the following parameters are estimated:

- Intercept and sex- and ICV-coefficient for mean initial latent volume

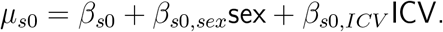

- Intercept and sex- and ICV-coefficient for mean initial change rate

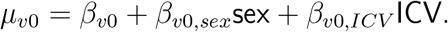

- Covariance matrix **Σ**_*s*0_ for initial latent variables.
- Age dependent bias *b*(*t*) for the acceleration parameterized as a cubic spline with 5 knots.
- Age dependent variance 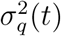 for the acceleration parameterized as a cubic spline with 5 knots.
- A bias for each site *β*(site).
- Feedback terms *f*_*s*_ and *f*_*v*_.
- Measurement noise variance for each site 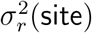.

When counting all the weights of the splines and the bias for each site, there is a total of 99 fitted parameters.

#### 2.2.3 Properties of the Stochastic Linear Dynamical System

The model can be expressed compactly in matrix form as:

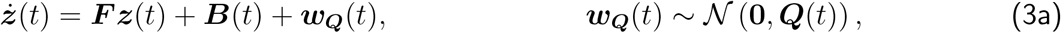

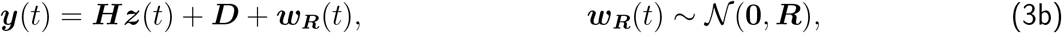

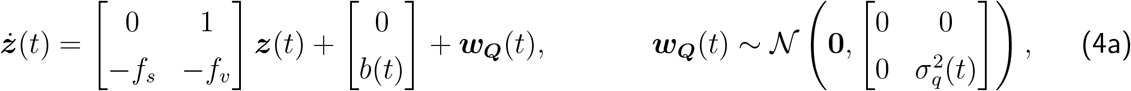

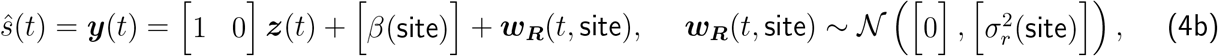

where ***y***(*t*) is the measurement vector which in our case is just the measured volume *ŝ*(*t*). A state-space diagram of the model is shown in Fig. 2.

**Figure 2:**
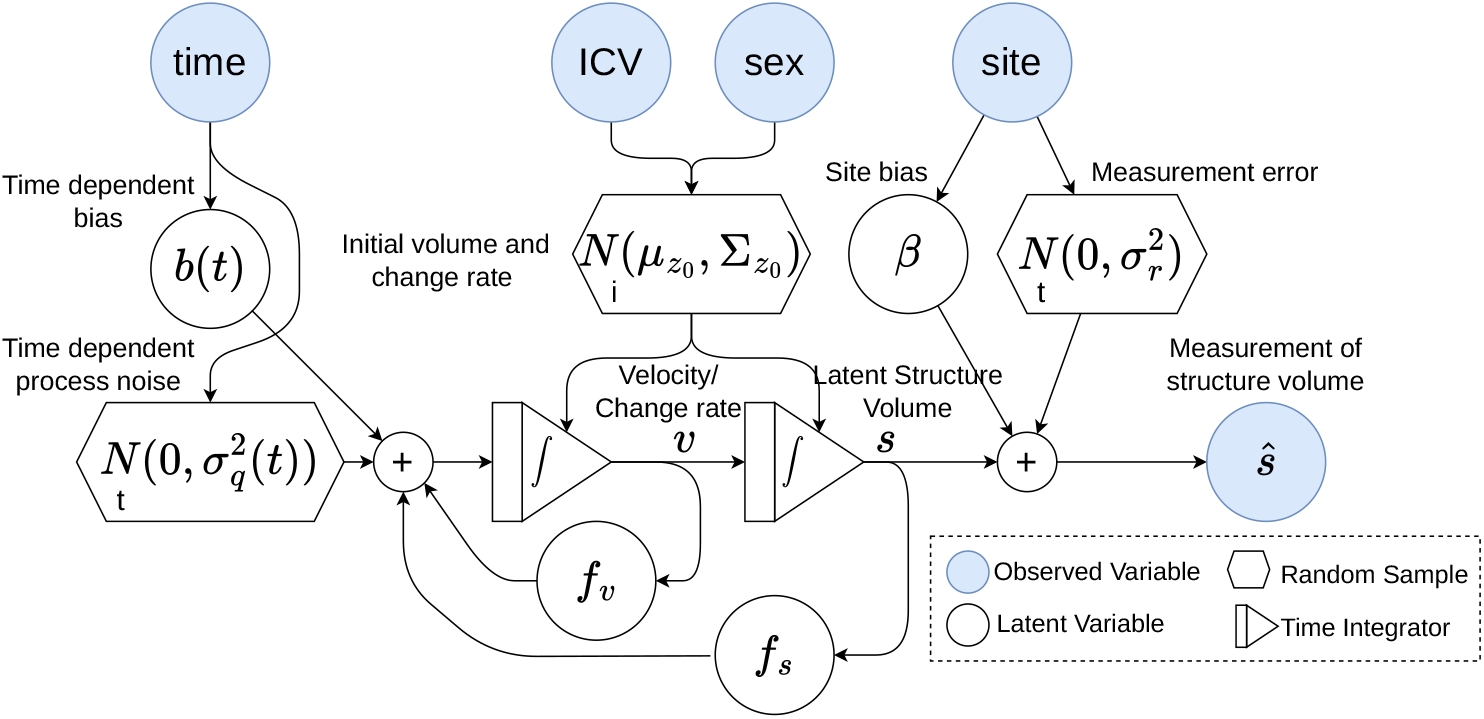
State-space diagram showing the dynamical model of the brain structure volume. The *i* in the hexagon indicates that the variable is iid for each individual, while a *t* indicates iid for each time step.

The continuous model can be time discretized analytically with a sampling period of Δ (Driver & Voelkle, 2018). We assume a zero-order-hold for the time dependent parameters *b*(*t*) and *σ*_*q*_(*t*) within each period Δ (Dahdah & Forbes, 2025; Simon, 2006), meaning these parameters remain constant within this period. Discrete variables are denoted with a star (∗), and time steps are indexed by *k*, such that ***F*** ^∗^ represent the time discrete version of ***F*** and *b*_*k*_ := *b*(*t*_0_ + *k*Δ). The time discrete model is given by

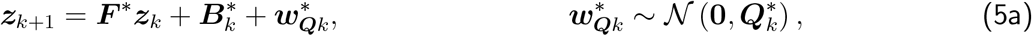

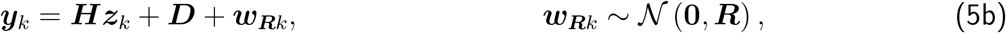

where the ***F*** ^∗^, 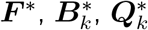 can by computed numerically from ***F***, ***B***(*t*) and ***Q***(*t*) using a matrix exponential (Dahdah & Forbes, 2025). The initial conditions remain as specified in Eq. 2c.

Assume that at time step *k*, the latent state is estimated as ***z***_*k*_ ~ 𝒩 (***µ***_*k*_, **Σ**_*k*_). ***µ***_*k*_ represents the state mean (which can be both the population norm or the mean of an individual’s posterior distribution after measurements) and **Σ**_*k*_ denotes the associated uncertainty. The distribution for the subsequent time step is given by

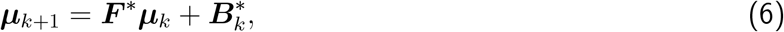

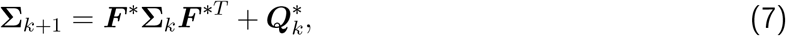

which for *n* time steps becomes:

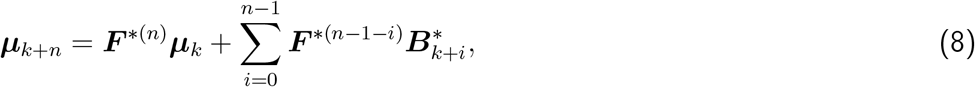

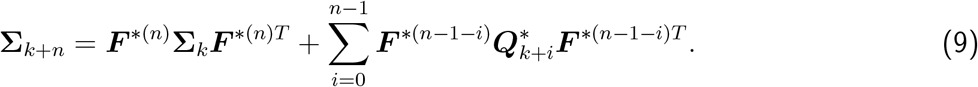

This model places an important constraint on the system by coupling the latent change rate with the volume over age and thus increases the statistical power of analyzing variability in change rate.

#### 2.2.4 Correlation Characteristics of the Stochastic Dynamical System

Our goal is to quantify the explained variance of brain structure volume in adulthood given the initial volume at young adulthood *t*_0_. To do so, we use the auto-correlation (AC) which is the normalized auto-covariance.

The auto-covariance ***K*** with lag n is defined as

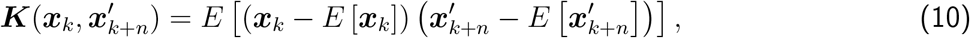

for the random variables ***x***_*k*_ and 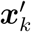. We will only consider the case where *n* ≥ 0, since the autocovariance is symmetric for negative *n* and this simplifies the equations a little. Let ***d***(**Σ**) be the vector of the diagonal of a matrix **Σ**. We define the standard deviation vector as 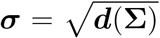 and the outer product of this vector as 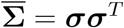. The auto-correlation ***ρ***_AC_ is given by

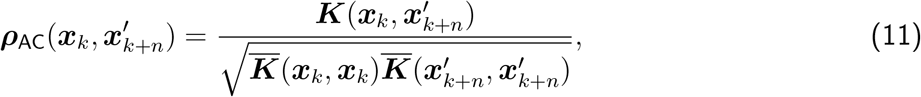

where the division is performed element-wise. Note that the auto-correlation ***ρ***_AC_ reduces to the correlation ***ρ*** when *n* = 0.

For a linear system such as Eq. 5, the auto-covariance has a closed-form solution given by propagating the initial covariance through the system transfer matrix ***F*** ^∗^ *n* times, which gives

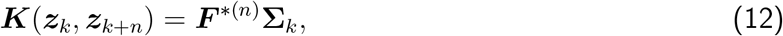

and

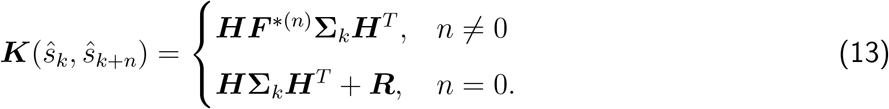

The measured volume has the same co-variance with the latent variables (including the latent volume) as the latent volume itself, which gives,

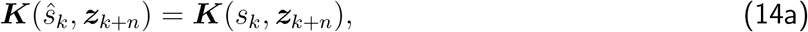

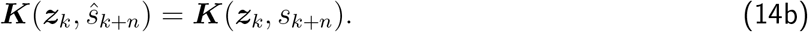

***ρ***_AC_ for the latent variables can be written compactly as

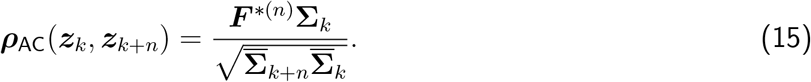

For the measured volume, the measurement noise must be added to the total variance. Thus, ***ρ***_AC_ for the measurements with *n >* 0 becomes

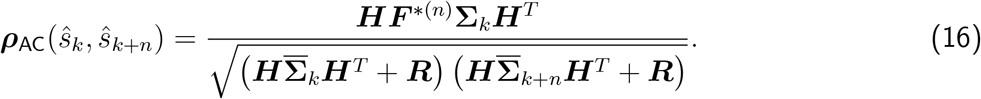

By using Eq. 14 one can obtain similar correlations between *ŝ*_*k*_ and ***z***_*k*+*n*_.

We assign names to the elements of Eq. 15 according to their interpretation as correlation coefficients:

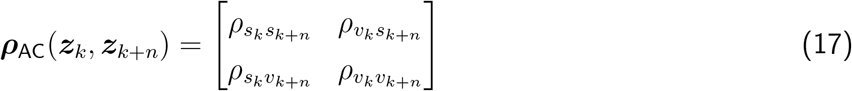

An intuitive explanation of these coefficients, assuming comparison between time points *k* and *k* + *n*, is as follows:

- 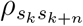 is the correlation coefficient of the structure volume at time step *k* and at the later time step *k* + *n*.
- 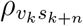 is the correlation coefficient of the change rate at time step *k* and the volume at the later time step *k* + *n*.
- 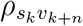 is the correlation coefficient of the volume at time step *k* and the change rate at the later time step *k* + *n*.
- 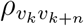 is the correlation coefficient of the change rate at time step *k* and at the later time step *k* + *n*.

#### 2.2.5 Estimates of Change Rate from Measurements

As the brain ages, the accumulated differences in anatomical changes affect the volume of the brain structures. Consequently, a single volume measurement, when compared to the age-group norm, can serve as an estimate of the current rate of change. However, earlier changes may not accurately reflect the current rate, and the accumulated changes might be too small compared to measurement noise for the volume to be related to the current change rate. To evaluate how well a single volume measurement of brain volume can act as a proxy for the current change rate, we compute the correlation between the measured volume at a given age and the latent change rate given by 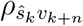.

An alternative method for obtaining an estimate of the change rate is to take the difference between two volume measurements, which we call a delta estimate and denote 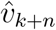. This is given by

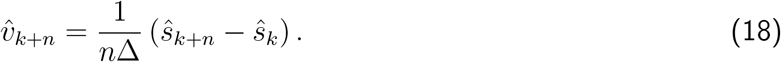

The question is whether this is a good measurement of the true change rate at the last time step *k* + *n* of the two measurements. We quantify this by calculating the correlation *ρ* between 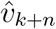 and *v*_*k*+*n*_ which is given by

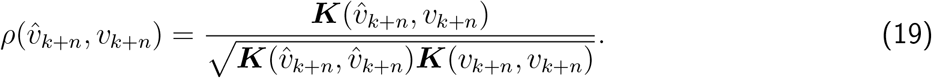

A little mathematical manipulation results in the following expression for the normalized correlation:

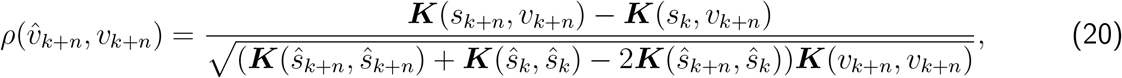

which are all quantities that can be calculated from Eq. 12 and Eq. 13.

Another common approach to estimate the change rate is to take many samples within a period of time, and use the slope of a linear model as the estimate. Let ***ŝ***_*k*:*k*+*n*_ = [*ŝ*_*k*_, *ŝ*_*k*+1_, …, *ŝ*_*k*+*n*_] be the vector of the equally spaced measurements. It can be shown that the slope *α*(***ŝ***_*k*:*k*+*n*_) is given by

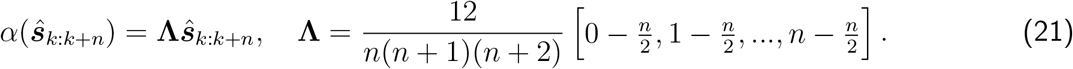

One can thus calculate the correlation as

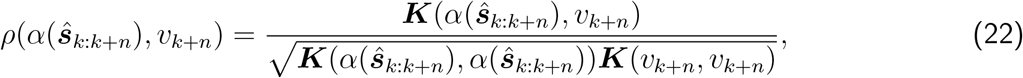

which again with a little mathematical manipulation can be shown to be

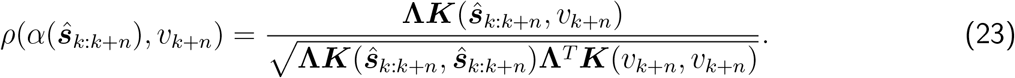

In Appendix A, similar derivations are shown for a linear model with a time-constant change rate. This provides analytical expressions for the standard deviation of the slope estimate and signal-to-noise ratio (SNR) with respect to the total scan interval, scanning strategy, and measurement noise.

#### 2.2.6 Implementation Details

The posterior distribution of the latent variables *p*(***z***_*k*_|*y*_1_…*y*_*K*_) for each individual can be calculated analytically using a Kalman/RTS smoother (Kalman, 1960; Murphy, 2023; Rauch et al., 1965).

Similarly, the exact model likelihood of the longitudinal observations,

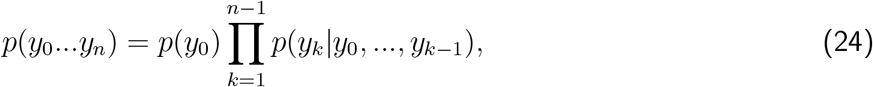

can be evaluated by marginalizing out ***z*** analytically. The model, therefore, naturally handles both a single and multiple observations by calculating the joint probability of all the observations.

We parameterize 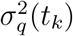 and *b*(*t*_*k*_) as cubic splines with 5 control points. The volume data is discretized to a sampling interval Δ of 1.0 year. We tested intervals of 0.5 years and 2 years, which yielded the same results. The priors of the variance terms are parameterized as a squared softplus (Dugas et al., 2000) transformed Gaussian distribution.

The model is implemented using the Python package Pyro (Bingham et al., 2019). Maximum a posteriori (MAP) estimates are first found using the Limited-memory Broyden-Fletcher-Goldfarb-Shanno (L-BFGS) algorithm (Nocedal & Wright, 2006). A Markov Chain Monte Carlo (MCMC) algorithm with a No-U-Turn (NUTS) sampler (Hoffman, Gelman, et al., 2014) is initialized to the MAP estimates, as recommended by Geyer, 2011, and 1000 samples from the posterior are drawn after a warm-up of 300 samples. The trace plots can be examined in the Supplementary Materials.

We used quite wide priors on the measurement errors of the structures derived from controlled experiments in Vidal-Piñeiro et al., 2025. That is, we centered the priors of *σ*_*r*_ around the estimated measurement standard error for each structure and set the prior standard deviation to the same number.

The combined sample has a high density of individuals in the age range of 50 to 70. To ensure that the analysis was invariant to the age distribution, we fitted the model to the data from only LCBC and Cam-CAN, which have a more even age distribution. This yielded the same general results as using the full dataset, but with greater uncertainty in the estimates. These results can be seen in Supplementary Materials Fig. 1 and 2.

## 3 Results

### 3.1 Dynamical Aging Norm Curves for Brain Structures

Fig. 3 shows the normative aging dynamics of brain structures across the adult lifespan. The mean and two standard deviations around the mean are plotted for both the latent volume and the change rate. The mean trajectories of the volumes show distinct shapes. The hippocampal and supratentorial volumes gradually decrease until around 55 years of age, followed by an acceleration of the rate of change and, therefore, faster shrinkage after age 60 years. The white matter and the brain stem volume follow similar mean trajectories, with an increase in volume up to the age of 50 years, followed by a decline. Gray matter volume shows a steady, close to linear, shrinkage throughout life. The lateral ventricles follow a very different trajectory. At the age of 18, the lateral ventricles are small; up to the age of 50 years, there is a small but measurable increase in volume. After that, the rate of growth accelerates, leading to a rapid increase in size.

**Figure 3:**
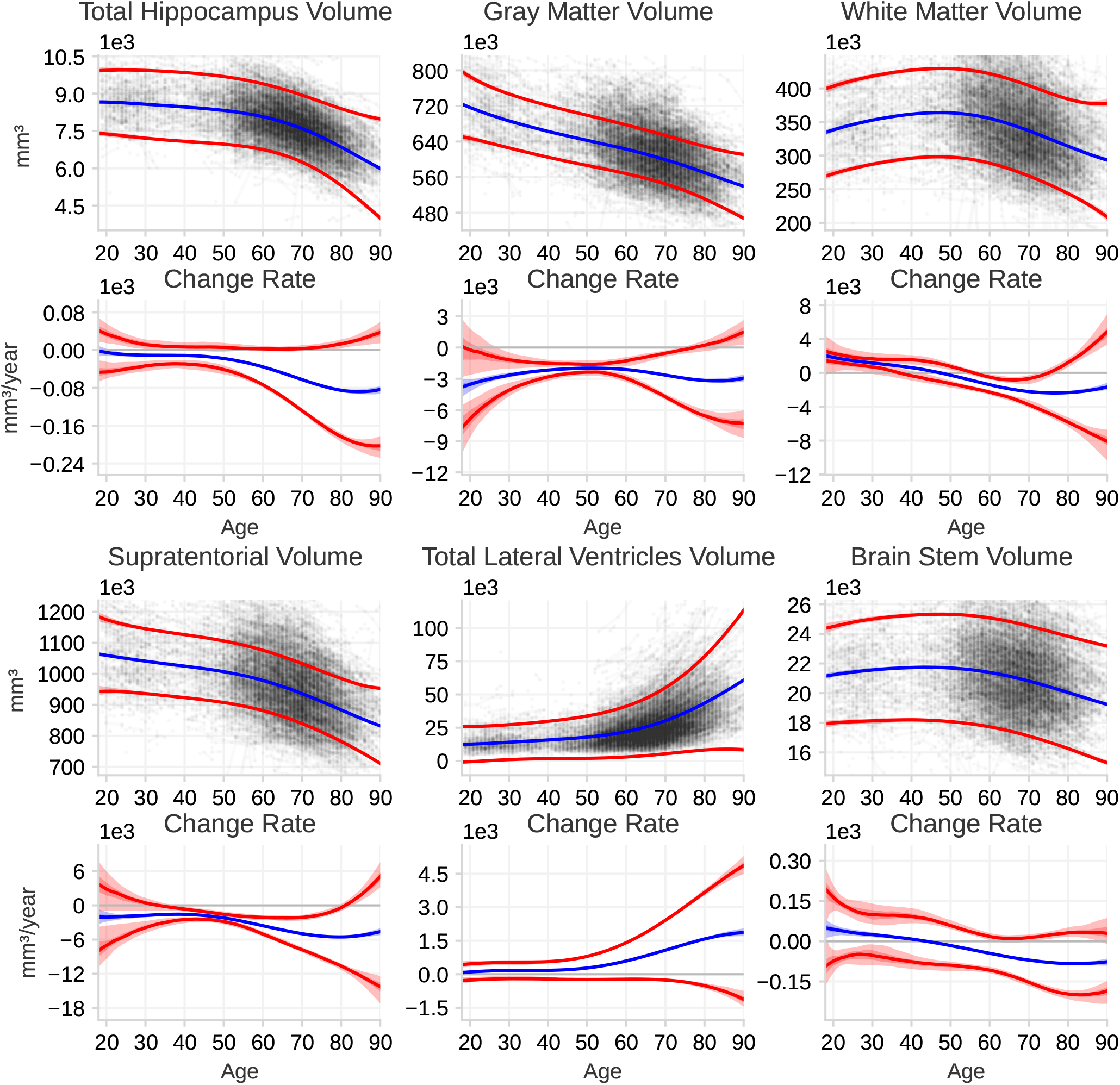
Norm curves for the dynamics of selected brain structures. For each structure the top plot shows the latent structure volume for the population from age 18 to 90 years, and the bottom plot the change rate of the latent volume. The observed data is plotted in the background. The blue curves indicates the population mean and the red curves show two standard deviations from the population mean, meaning ~95% of the population is expected to have volumes and change rates within this region. The shaded region indicate the 50% and the 95% confidence interval (CI) for each curve. For some curves the Cis are small, and therefore the CI regions appears as a single line.

The variation in individual change rates for the hippocampus, total gray matter, and supratentorial volume follows similar age-related patterns. Between ages 18 and 30 years, our data provide insufficient statistical power to precisely quantify how much individuals differ in their change rates. However, by around age 30 the estimates indicate that inter-individual variation in change rate is small. From approximately 30 to 60 years, variation in change rate remains low. After about age 60, variation increases markedly, resulting in large inter-individual differences in change rate by age 80. White matter volume shows small variations in inter-individual change rates up to the age of 60 and increases thereafter, while the brain-stem has small but stable inter-individual differences in the change rate throughout life. The ventricles show a steady variation in the inter-individual change rate up to the age of 50 and then drastically increase afterwards. For comparison, at age 70, the ratio of the inter-individual differences in change rate to the mean volume is approximately 6 times larger for the ventricles than for the hippocampus.

### 3.2 Explained Variance from Young Adult Levels vs. Systematic Change

The previous section shows that by the age of 30 years, the brain structure volumes have settled at a stable level. We estimated how much of the variance in later-life brain-structure volume can be accounted for by volume level at age 30. Using the fitted model, we derived the *R*^2^ as the squared correlation between volumes at age 30 and at later ages, after accounting for intracranial volume (ICV) and sex so as to maximize sensitivity to change. The proportion of explained variance is shown in Fig. 4. We computed *R*^2^ both in the absence and presence of measurement noise. Without noise, it indexes how much variance in *latent* later-life volume is explained by sex, ICV, and *latent* volume at age 30; with noise, it indexes the variance in *measured* later-life volume explained by sex, ICV, and *measured* volume at age 30.

**Figure 4:**
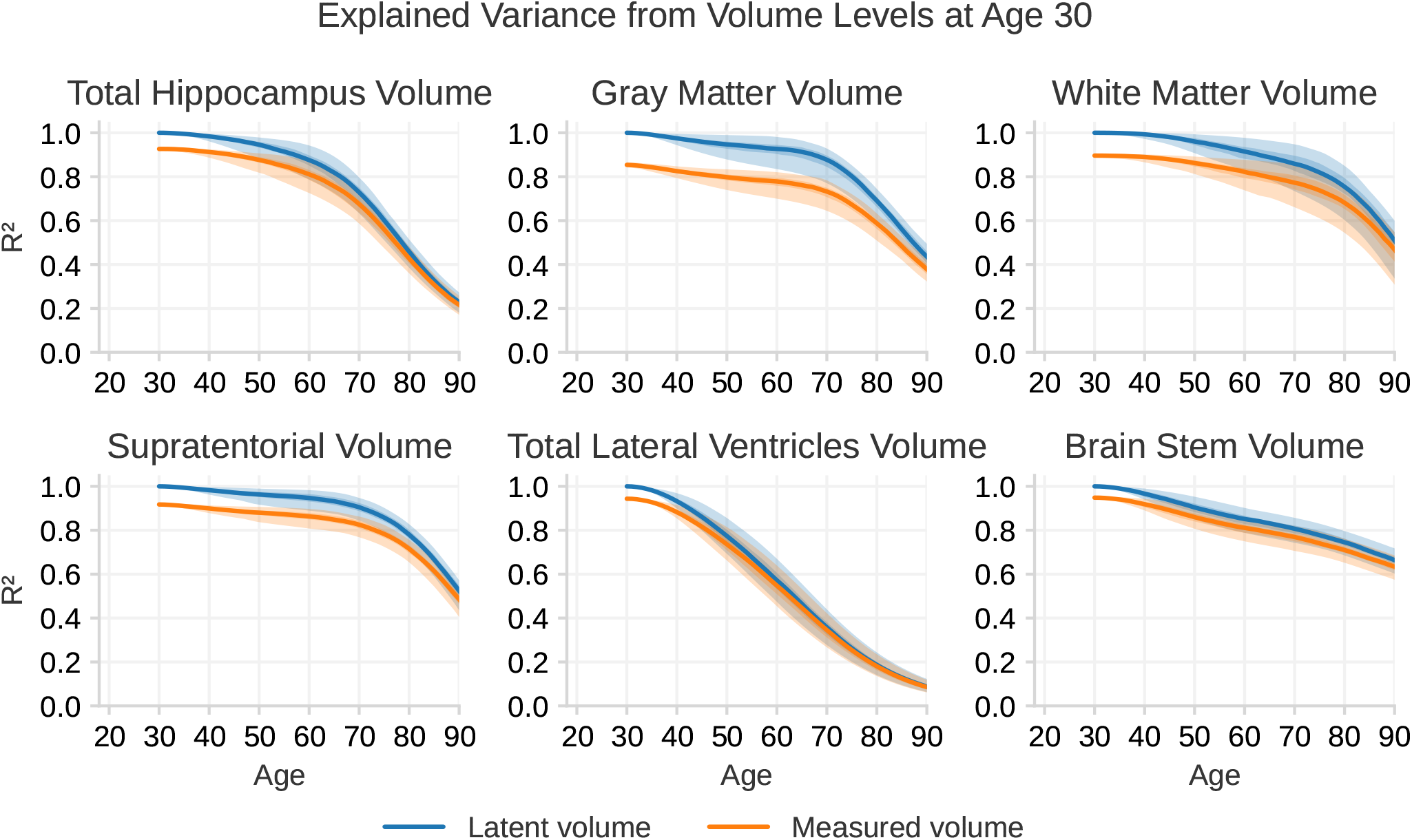
The ratio of explained variance *R*^2^ in brain structure volumes across the lifespan, given the structure volume at age 30 years. The blue line shows the explained variance of the latent volumes without measurement noise, while the orange line includes measurement noise. Both lines are calculated from the fitted model. 95% Confidence intervals are shown as shaded regions around each line; however, in most graphs, the intervals are narrow and appear as lines rather than shaded areas. In this plot 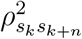 and 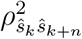 with *t*_*k*_ = 18 years of Eq. 17 are visualized.

We observe that, up to the age of 60 years, the stable volume level established at age 30 explains nearly all of the variance. For instance, for hippocampal volume at age 60 years, the model predicts that 87.39% (CI 78.8–94.2%) of the variance is explained by the early-adulthood level, whereas the corresponding measured volume has 80.7% (CI 72.6–87.13%) of its variance explained by a volume measure at the young-adult level. This value then drops rapidly with the large increase in systematic differences in volumetric change, so that by age 80, only 45% (CI(38.6–51.4%) of the variance is explained. The other structures show similar trends. The exception is again the lateral ventricles, which show a steady decrease in explained variance from the initial volume throughout the age range; i.e., individual differences in change are observed at earlier ages. For example, the explained variance in ventricular volume at age 50 years attributable to the initial volume is 77.6% (CI 69.3–85.5%), and this drops steadily so that by age 90 years the initial volume accounts for only 8.91% (CI 6.31–12.3%) of the variance.

### 3.3 Estimation of Change Rate

Next, we examine how well one can estimate the individual volumetric change of a brain structure. We examine three methods for obtaining an estimate of change: i) Using a single measurement of the volume and comparing that to the norm, accounting for age, ICV, and sex. Volumes deviating from the norm are likely to have experienced and continue to experience individualized change. ii) Using the time-normalized difference in volume between two measurements to provide a delta estimate of the change rate. iii) Obtaining more than two measurements and using the linear regression slope as an estimate of the change rate.

The correlations of the volume with the change rate are shown as black lines in Fig. 5. From our analysis, we see that up to the age of around 65 the correlation between a volume measurement and the latent change rate is either very low or even negative. For instance, at the age of 50 years, the measure of hippocampal volume has a correlation of −0.17 with the change rate at that age. The negative sign is due to the hippocampus shrinking, and larger structures are shrinking at a higher rate. After a few years of accumulated individualized changes, the correlation between volume and change rate increases, leading to a positive correlation at later ages. For instance, the correlation between the measured volume and latent change rate is around 0.33 at age 80. The lateral ventricles have, in contrast to the other structures, by age 40, a significant positive correlation of 0.35, which increases steadily to 0.65 at the age of 80 years.

**Figure 5:**
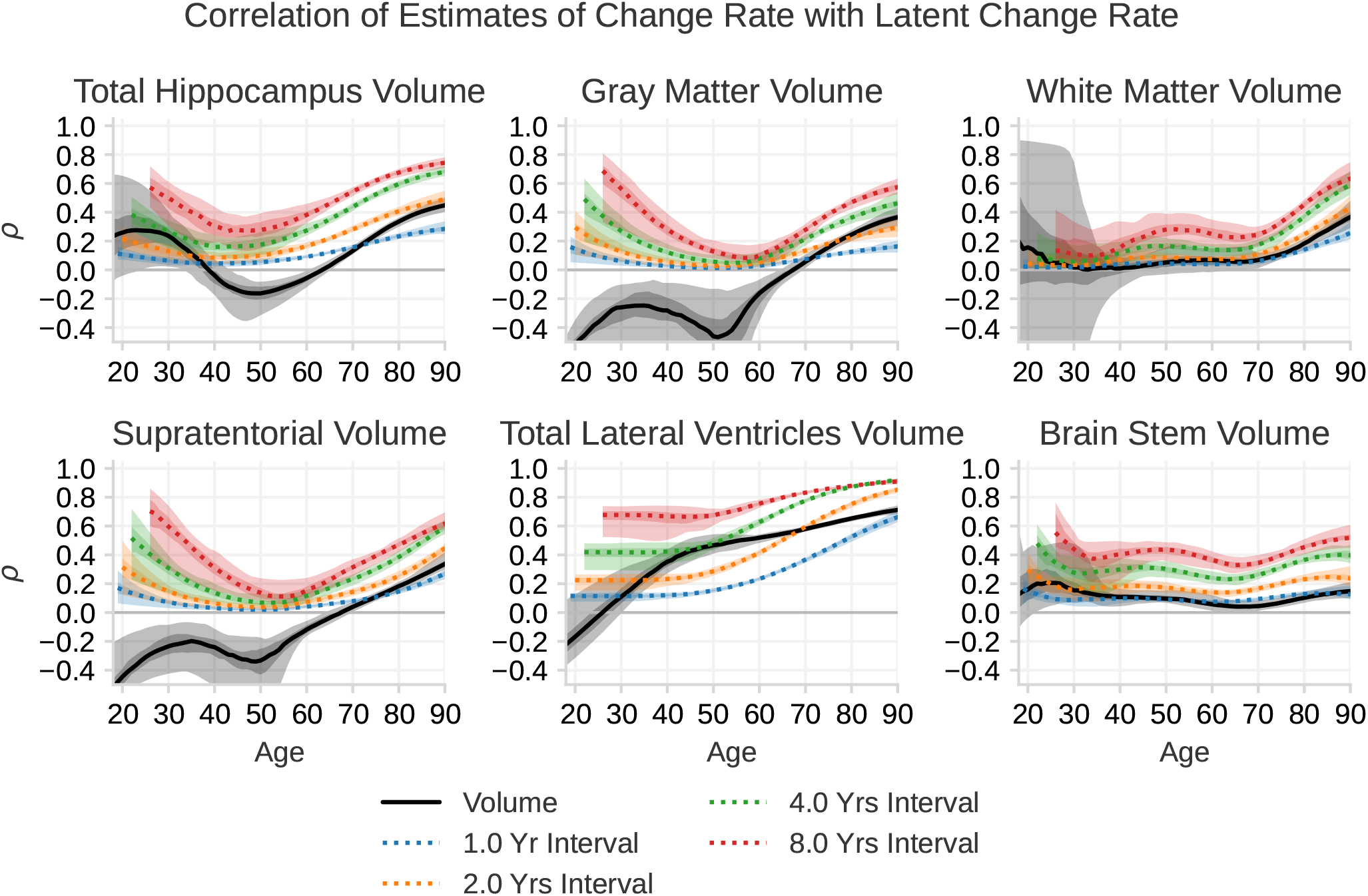
The correlation *ρ* between measured volume with the latent change rate, and the correlation of the delta estimate with latent change rate. The black line indicate how much a measurement of the volume of a brain structure correlates with the latent change rate of that volume at that time point. The colored lines indicates how much the the slope between two longitudinal measurements of brain volume correlates with the latent change rate of that volume at the last session. Each color indicates the time between the sessions. The shaded regions show the 95% and 50% confidence interval of each line. The black lines correspond to 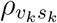 of Eq. 17 with *n* = 0 and the colored lines correspond to 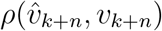 of Eq. 20.

Our analysis of the delta estimate is shown as colored lines in Fig. 5. We find that estimating individual change from delta estimates results in a correlation with a U-shaped curve over age for hippocampal volume, gray matter volume, and supratentorial volume. However, due to the lack of statistical power at the youngest ages, we cannot confidently say whether measuring change at this age is possible. In the middle of the lifespan, we can confidently say that the correlation is very low. For instance, for gray matter volume, the correlation is only around 0.1 at age 50. With an increase in the variability of the change rate at older ages, a delta estimate of change is again able to detect individualized change, but only with modest correlations. For instance, the correlation with true change at 80 years and a 2 year follow-up for gray matter is 0.24. For white matter and brain stem volume, we find low correlation with true change throughout life, with a slight increase for white matter at the age of 80. The ventricles show the largest and most stable correlation with true change from delta estimates, which, already at a young age, shows a similar correlation with latent change as gray matter does at an older age. Interestingly, we find that performing a delta estimate within one year has a lower correlation with true change compared to a volume estimate after the age of 70 for most structures.

Fig. 6 shows the correlation between estimates of change and true change when using multiple sessions. We observe that increasing the number of sessions only shows a moderate increase in correlation compared to increasing the time between the first and the last session. For instance, for a 2-year measurement interval for hippocampal volume at age 80 years, the correlation with the change rate is 0.43 using 2 measurements and 0.54 using 12 evenly spaced measurements within the same scanning interval. This improvement is lower than for going from 2 to 4 year intervals with only two scans, where the correlation improves from 0.43 to 0.62.

**Figure 6:**
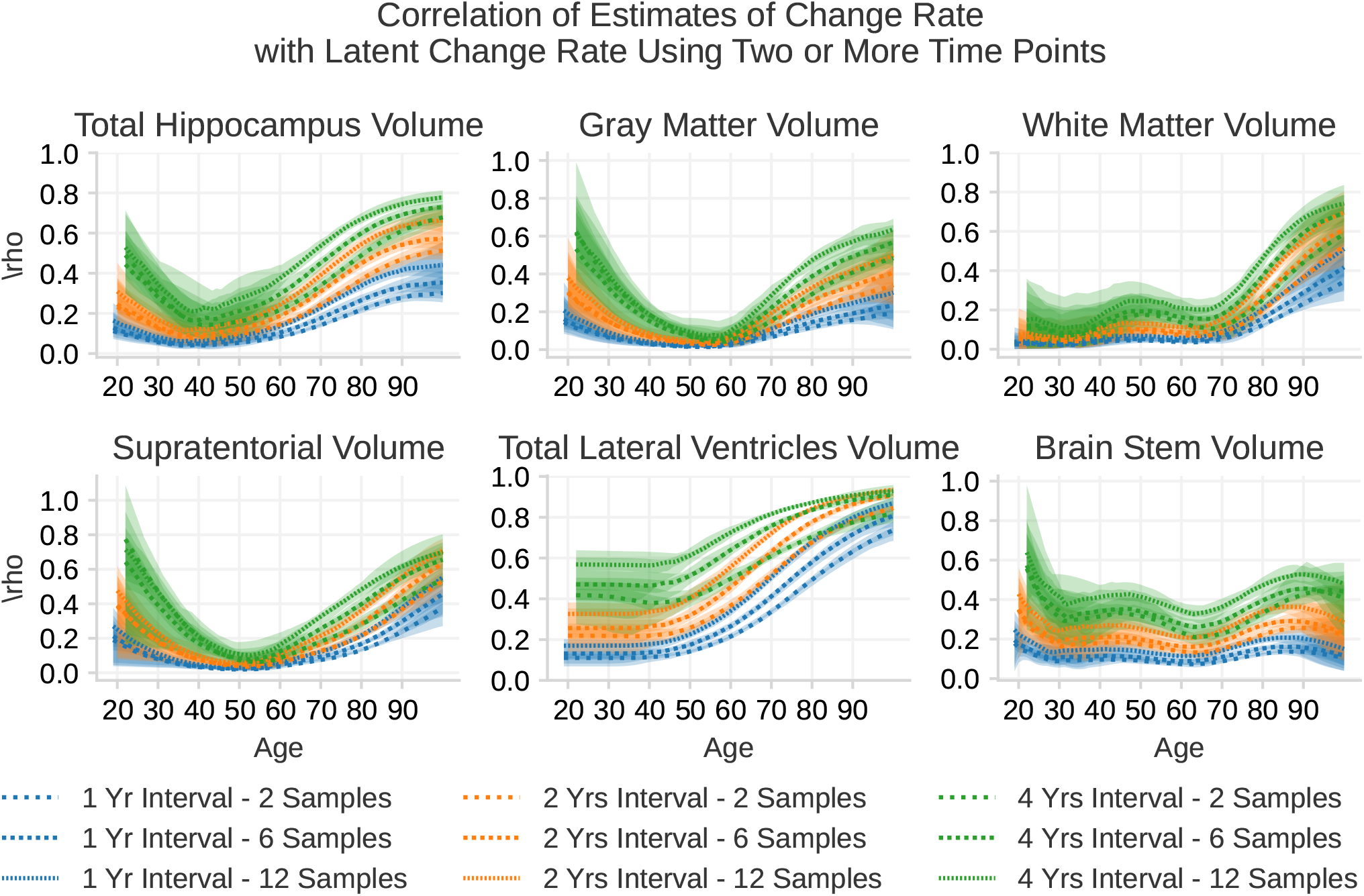
The correlation *ρ* between slope estimates of change rate from 2, 6 and 12 sessions with the latent change rate. The 2-samples lines are the same as the colored lines in Fig. 5 while the other lines indicate the correlation between the slope estimate with the latent change rate when including more intermediate sessions. These correlations are calculated from Eq. 23.

From our calculations, increasing the number of sessions from 2 to 12 is almost never as good as doubling the time between two measurements. The take-home message is that increasing the signal, i.e. the absolute amount of change, is more efficient than adding more measurements. Note that these calculations assume equal spacing between each measurement, and the situation will be different if multiple measurements are performed close to the first and the last time point. See Appendix A for an exploration of how the sampling distribution affects the sensitivity to change.

## 4 Discussion

The present analysis yielded three main results. First, we find that the systematic inter-individual differences in volumetric brain change before age 60 years are much smaller than the differences seen later in life in a large cross-cohort sample of cognitively healthy adults. This indicates that in cognitively healthy adults, morphometric differences prior to age 60 years largely reflect stable individual differences, and are for the most part not ascribed to differences in change rate. From about 60 years of age, individuals began to follow detectably different trajectories, and differences in change gradually explained more variance with increasing age. Second, because the variation appears to be dominated by stable individual differences before the age of 60, a model trained to predict cross-sectional age will be sensitive to age, but not differences in aging. The only measure partially reflecting individual differences in change before age 60 was the ventricles, which are known to be sensitive to age-related changes, but typically do not dominate in brain age calculations. Finally, the reliability to detect differences in change was very low in the age range 30 to 60 years, even with many samples and up to 8 years between first and last measurement. From the age of 60 years, reliability increases with longer follow-up times having the largest effects, while adding more intermediate time points comparatively affected reliability to a very modest degree. For example, two scans 4 years apart yielded similar or higher reliability to detect change than 12 scans evenly distributed over 2 years. The implications of the findings are discussed below.

### 4.1 Cross-sectional Differences Mostly do Not Reflect Differences in Brain Change

The aging norm trajectories of the different brain structures replicate previous aging studies (Fjell et al., 2013). All the structures showed age-related decline, but followed distinct patterns. The hippocampus remains relatively stable until around 60 years of age, after that accelerated decline is observed. In contrast, total gray matter volume exhibits almost linear reductions, with a slight acceleration from age 60 onward. White matter and brain stem are the only structures that do not show a monotonic decline; instead, they apparently increase until middle age, which is a well-established pattern (Allen et al., 2005; Walhovd et al., 2005). The lateral ventricles display the most unique trajectory, with a small increase in volume until around 50 years, followed by a rapid increase throughout the rest of the lifespan.

Importantly, as was recently demonstrated in a study of the UK Biobank database, high cross-sectional age correlations do not necessarily translate to high sensitivity to brain change (Smith et al., 2025). This means that if individuals have a smaller volume of an age-sensitive brain structure, this does not imply that they have undergone greater age-related changes than others. The present results clearly confirm that while brain regions may be age-sensitive at the group level, this does not reflect the existence of systematic differences in age-related rates of change (Vidal-Piñeiro et al., 2025). We observe substantial agerelated changes in all tested neuroanatomical volumes, with some, particularly total gray matter volume and supratentorial volume, changing continuously from early adulthood. Nevertheless, the variance attributable to young adult levels surpasses the variance associated with individual differences in change by far, at least until advanced ages.

As shown in Figure 4, until around 60 years of age, nearly all differences in neuroanatomical volumes reflect stable, individual differences established in early adulthood. From approximately 50 years, systematic differences in change rates between participants increases. These differences accumulate over time, and at 60 years, their impact is sufficiently large to be observed as smaller volumes, but only a minor proportion of individual variation is still due to systematic differences in change. This proportion steadily increases at different rates across structures, reaching approximately 10-20% at 70 years and 50% or more at 90 years for most structures. It is crucial to note that these findings represent group-level effects and cannot be directly translated into considerations on level versus systematic change in the brains of individual participants.

The reason for the small proportion of variance attributed to systematic differences in change is that absolute volumetric differences between individuals are substantial, while inter-individual variation in change is relatively small. Consequently, only after several decades do volumetric differences significantly reflect change. However, the ventricles deviated from this pattern, and we are able to detect small individual changes throughout the first 30 years of adulthood before expanding rapidly from around 50 years. The relatively minor individual differences in baseline volume, combined with systematic variation in individual change rates, also at young age, result in a situation where a much larger proportion of volume variation can be attributed to change. Even at 50 years, around a quarter of the volumetric differences between individuals can be explained by differences in change, a proportion that reaches 90% at 90 years. This means that ventricular size is the only tested brain measure that reflects actual agerelated variability as a result of accumulated individual changes when measured at a single time point throughout life. This observation was previously reported by Vidal-Pineiro et al., 2021.

Thus, measures where the rate of change later in life is large relative to inter-individual variation in young adulthood will, by this reasoning, reflect proportionally more change than baseline effects when measured in aging. Another good example is white matter hypointensities, not measured here, where variation appears to reflect change to a greater extent than most other brain measures (Smith et al., 2025). However, this also means that, for other brain measures, cross-sectional comparisons do not inform us, for the most part, about individual differences in aging in cognitively healthy normal populations. Hence, typical brain age models trained to predict chronological age will typically be dominated by crosssectional variance and, consequently, are very insensitive to individual differences in the rate of aging (Smith et al., 2025).

### 4.2 Impact of Levels vs. Change will Depend on the Sample

An important consideration is that the proportion of variance at a given age that can be attributed to levels at young adulthood vs. systematic differences in change will depend on the type of samples included. For example, we find that about 50% of the individual differences in hippocampal volume at age 80 years can be ascribed to differences in change, but this number is likely much higher in neurodegenerative disorders like AD. While we estimate annual change in hippocampal volume to be around 1% after 60 years, this number often exceeds 3.5% in AD patients (Fjell et al., 2009). Consequently, such a large difference in rate of atrophy will eventually manifest in lower hippocampal volume in AD patients, regardless of the variation at young adulthood. Although large baseline hippocampal volume may be a protective factor, volumetric differences between patients with progressed AD and age-matched controls will likely to a larger extent reflect accumulated higher atrophy rates in the patients. How much of the variance that can be ascribed to each remains to be tested. Similarly, comparing other patient groups having conditions associated with magnified volumetric change rates may also be expected to yield a higher proportion of volumetric differences ascribed to change.

Still, AD and some other age-related dementias occupy the high end of the neurodegeneration spectrum. For many other conditions where neuroanatomical differences are found — such as neuropsychiatric disorders like schizophrenia (Constantinides et al., 2023; Van Erp et al., 2018; Zhu et al., 2023) or major depression (L. K. Han et al., 2021), differences in change rates are usually much smaller (Abé et al., 2022; Vita et al., 2012) than in AD. In such cases, the present results suggest that volumetric differences largely reflect stable levels, likely originating early in life and possibly partly due to developmental differences.

For example, a large multi-sample study of bipolar disorder did not find substantial differences in change rates of neuroanatomical volume between patients and controls (Abé et al., 2022). Therefore, a crosssectional comparison would be expected to reveal primarily level differences between groups, regardless of the age at testing. However, this study included several middle-aged and older participants and higher ventricular expansion was observed in the patient group. This finding is particularly relevant, as ventricular expansion was the one variable in the present study that reflected the most systematic differences in change.

Other cross-sectional studies of schizophrenia have found that patients have higher predicted brain age than controls (Constantinides et al., 2023). As argued above, brain age prediction models depend on age relationships and, therefore, primarily reflect the features of brain structures that are most reliably linked to age. For example, total gray matter volume would typically be assigned high weight in brain age models, and has been reported to be the measure most predictive of brain age differences between patients with schizophrenia and controls (Ballester et al., 2023). However, as low inter-individual differences in change yield more precise brain age predictions, such measures are likely insensitive to aging (Smith et al., 2025). An implication is that cross-sectional studies of aging differences between neuropsychiatric groups run a high risk of confounding stable levels with true aging effects.

Education, sometimes regarded as one of the strongest protective factors for dementia (Livingston et al., 2024), has recently been shown to correlate with larger brain volumes but not with differences in brain change (Cox et al., 2016; Fjell et al., 2025; Nyberg et al., 2021). Longitudinal and cross-sectional brain structural measures are also found to relate differently to cognitive function (Walhovd et al., 2022). These results fit well with the present observation that differences in regional brain volumes mostly reflect stable individual differences, also in relatively advanced ages.

Conversely, when comparing groups with different clearly defined developmental risk factors, such as prenatal drug or alcohol exposure, very low birth weight, or premature birth, brain volumetric differences will probably for even longer reflect effects present early in life. Consistent with this, a review showed that reported effect sizes on neuroanatomical volumes from fetal life risk factors were several times larger than those from later-life risk factors such as hypertension, overweight, and heavy alcohol consumption (Walhovd et al., 2023). We expect the same to be the case with neurodevelopmental psychiatric diagnoses associated with brain morphometric differences, such as attention deficit hyperactivity disorder (Hoogman et al., 2017). Since these differences also originate from early in life, adult comparisons will be expected primarily to reflect stable levels even in old age (Walhovd et al., 2012).

### 4.3 Implications for Studies of Brain Aging

The results confirm a growing body of evidence suggesting that deviation from “age-expected” lifespan brain trajectories cannot be used to make inferences about brain aging, at least not before the age of 60. Cross-sectional studies can provide important information about the brain basis for cognitive function in aging, but brain aging per se is better studied using longitudinal designs. Interestingly, the results suggest that we have very little signal to detect systematic differences in brain change before the age of 60 years, although this will ultimately depend on the reliability of the measurements (signal-noise ratio) (Brandmaier, Wenger, et al., 2018; Narayanan et al., 2020; Vidal-Piñeiro et al., 2025) and the sample in question.

As has been shown previously using simulations and formal analysis (Brandmaier, Von Oertzen, et al., 2018; Brandmaier et al., 2024; Ghisletta et al., 2020) and computations based on empirical data (Hertzog et al., 2008; Vidal-Piñeiro et al., 2025), the results show that time between measurements is much more important for reliability than the number of observations. Except for the lateral ventricles, reliability was low even with 12 scanning session evenly spaced across 12 months. In contrast, a 4 year follow-up interval increased reliability greatly. This demonstrates that more signal is the most important factor for reliable measures of longitudinal brain change. Since change rates increase with age, the reliability also increased greatly with aging, reaching a correlation of about 0.70 for hippocampal change over four years even with only two scans. Hence, an implication for aging studies is to maximize follow-up time even at the expense of more time points.

An exception to this conclusion is, of course, when special events are of interest, such as in intervention and experimental studies, which differ from observational studies of naturally occurring aging processes. For example, intervention studies have detected systematic effects of cognitive training over a few weeks in younger and older participants (Bråthen et al., 2022; Engvig et al., 2010; Wenger et al., 2012, 2017). In these cases, post-intervention differences reflect change due to the experimental intervention. Another reason to prefer many imaging sessions is that participants are likely to drop out of the study, and acquiring intermediate sessions might be necessary to ensure some data before participants potentially drop out. In addition, some theoretical models of experience-dependent plasticity predict curvilinear changes in brain volume as a function of skill acquisition, in the sense that an early increase is followed by a decrease (Lövdén et al., 2020). Clearly, testing such a model requires more than two measurement occasions.

Another implication is that if only cross-sectional data are available, aging studies should focus on brain measures reflecting differences in change to a larger degree, e.g. the lateral ventricles and WM hypointensities (Smith et al., 2025; Vidal-Pineiro et al., 2021). Similarly, residualizing total brain volumes with respect to ICV has been shown to correlate with ongoing atrophy (Fürtjes et al., 2025), because these measures are very highly correlated in adolescence and young adulthood, and residual scores therefore partially reflect accumulated change since lifetime peak brain size. In any case, the results underscore the need to consider early-life developmental processes to understand and account for individual differences in brain structure at all later stages in life (Walhovd et al., 2016).

This study demonstrates the importance of improving the reliability of structural neuroanatomical measurements. Although sensitivity to change increases approximately linearly with the interval between sessions, waiting many years is impractical. As shown in Appendix A, 22 equally spaced sessions are needed to achieve the same reliability as simply doubling the time span; this approach is often impractical. Concentrating measurements at the endpoints (e.g., cluster scanning (Elliott et al., 2024)) reduces the theoretical requirement to 8 sessions to match that gain. However, this assumes truly independent scans acquired over a short period, which is empirically not the case (Elliott et al., 2024).

The volumetric measures analyzed here result from a chain of procedures, each contributing uncertainty to the final estimate, such as subject motion and positioning during scanning, MRI acquisition noise and scanner drift, as well as the uncertainty introduced by the segmentation and processing software. While it is difficult to identify which component is most tractable to improve, our analysis indicates that any reduction in the variance of the volume estimate leads to a linear improvement in the change estimate and is therefore of substantial practical importance. Thus, reduced noise with improved acquisition protocols would likely lead to more accurate detection of longitudinal changes.

### 4.4 Considerations and Limitations

Our results depend on the validity of the model assumptions. First, we assume that the measurement noise is Gaussian distributed and independent of all variables, such as age and structure size. This may not hold, as older individuals can be harder to scan due to factors such as movement while scanning (Vidal-Piñeiro et al., 2025). Second, we assume that the noise driving the stochastic component of the acceleration is also Gaussian distributed. The true distribution of change rate and acceleration is likely skewed, as shown in Vidal-Piñeiro et al., 2025. Third, similar concerns apply to the initial distributions of structural volume and change rate, which we also assumed to be Gaussian distributed. The model calibration curves, provided in the Supplementary Materials, indicate that the model is well calibrated for most structures, with the lateral ventricles showing the largest deviation from optimal calibration. This suggests that the Gaussian assumptions generally hold; however, ventricular volume could be modeled more accurately by incorporating higher moments.

There are also good reasons to assume Gaussian distributions. First, this choice makes the model analytical, so the probability of multiple observations from a single individual, as well as all the results presented in Fig 3 to 6, can be computed directly from the model parameters. Second, Gaussian distributions can be viewed as a natural baseline assumption. Most related models, such as Latent Growth Curve Models (Preacher, 2008) and the model used by Smith et al., 2025, also rely on Gaussian distributions.

It is important to consider that we assume participants observed at a given age are representative of the broader population at that age. In other words, we assume that individuals scanned in their 20s are a reasonable proxy for how individuals scanned in their 80s would have looked in their 20s (and vice versa), conditional on ICV. Including ICV may partially mitigate cohort-related differences that arise from developmental influences, given that ICV is established during development and remains stable in adulthood. However, this adjustment is unlikely to fully eliminate cohort effects, and will likely not eliminate period effects. Separating age-related change from period- and cohort-related differences is inherently difficult, particularly in pooled multi-cohort datasets, and residual cohort effects may therefore remain in our estimates (Rohrer, 2025). In addition, because cognitively impaired individuals were excluded, estimates, particularly at older ages, may be biased. In late life, when impairment is more prevalent, this exclusion likely yields a healthier-than-average sample and could lead to an underestimation of variability in rates of change.

Site-specific bias is modeled as an additive offset, which we treat as a pragmatic first-order approximation to site effects. A more flexible specification would include both additive and multiplicative terms, as in Beer et al., 2020, but this would further increase the complexity of an already complex model. In the Supplementary Materials, we show that the adopted correction substantially reduces site-dependent differences in the volumetric measures.

We used five knots for the splines, spaced equally over the interval from 18 to 100 years. As a result, changes in the dynamics that occur on shorter timescales are not captured by the model parameters. Fitting dynamical models with feedback and time-dependent parameters is challenging because the posterior over parameters exhibits strong dependencies. We therefore limited the splines to five knots to ensure posterior convergence.

### 4.5 Conclusion and Further Research

The present results show that by 30 years of age, structures in the brain have settled at a stable volume level that explains most of the variance up to the age of 60 years. After age 60, larger systematic differences in changes begin to accumulate, resulting in a gradual increase in brain neuroanatomical differences and in observed volumetric differences. Therefore, before the age of 60 years, differences in brain volumes primarily reflect stable levels from young adulthood. The proportion of variance attributable to systematic differences in change increases at higher ages, typically reaching 20-50% at age 80 years and 40-70% at age 90 years. The exception is ventricular volume, with more than 60% of the individual differences in volume at age 70 years reflecting differences in change, a proportion increasing to 90% at age 80. Further research should extend these findings to include different populations, including neuropsychiatric conditions, groups with different risk factors, and diseases known to affect brain structure. Clarifying the relative contributions of young-adult baseline levels and subsequent changes due to diseases and clinical conditions will be an important direction for future research.

## Ethics

Each contributing dataset was approved by its local Research Ethics Committee, and data were shared with us under the terms specified by the respective data-use agreements. Analyses were performed on secure institutional servers with restricted access, and no attempt was made to re-identify participants.

## Acknowledgments

This work was supported by South-Eastern Norway Regional Health Authority (to *EG* [HSØ-2021079]), the Department of Psychology, University of Oslo (to *KW, AF*), the Norwegian Research Council (to *KW, AM, DP* [ES694407]) and the project has received funding from the European Research Council’s Starting Grant scheme (to *AF* [283634, 725025] and *KW* [313440]). *AP-L* is partly supported by the Healthy Aging Initiative, the Eleanor and Herbert Bearak Memory Wellness for Life Program and grants from the National Institutes of Health, Jack Satter Foundation, and BrightFocus Foundation.

Each contributing dataset is supported by different sources. LCBC: the Norwegian Research Council (to *AF, KW*), and the National Association for Public Health’s dementia research program (*AF*). Umeå (BETULA): a scholar grant from the Knut and Alice Wallenberg (KAW) foundation to *LN*. UB (Barcelona): *DB-F* was funded by an ICREA Academia Award. *DB-F* acknowledges the CERCA Programme/Generalitat de Catalunya and is supported by María de Maeztu Unit of Excellence (Institute of Neurosciences, University of Barcelona) MDM-2017-0729, Ministry of Science, Innovation and Universities. *LW* and data collection in COGNORM is funded by the South-Eastern Norway Regional Health Authorities [#2017095] The Norwegian Health Association [#19536] and by Wellcome Leap’s Dynamic Resilience Program (jointly funded by Temasek Trust) [#104617]. The funding sources had no role in the study design.

Parts of the data collection and sharing for this project were funded by the Alzheimer’s Disease Neuroimaging Initiative (ADNI) (National Institutes of Health Grant U01 AG024904) and DOD ADNI (Department of Defense award number W81XWH-12-2-0012). ADNI is funded by the National Institute on Aging, the National Institute of Biomedical Imaging and Bioengineering, and through generous contributions from the following: AbbVie, Alzheimer’s Association; Alzheimer’s Drug Discovery Foundation; Araclon Biotech; BioClinica, Inc.;Biogen; Bristol-Myers Squibb Company; CereSpir, Inc.; Cogstate; Eisai Inc.; Elan Pharmaceuticals, Inc.; Eli Lilly and Company; EuroImmun; F. Hoffmann-La Roche Ltd and its affiliated company Genentech, Inc.; Fujirebio; GE Healthcare; IXICO Ltd.; Janssen Alzheimer Immunotherapy Research & Development, LLC.; Johnson & Johnson Pharmaceutical Research & Development LLC.; Lumosity; Lundbeck; Merck & Co., Inc.; Meso Scale Diagnostics, LLC.; NeuroRx Research; Neurotrack Technologies; Novartis Pharmaceuticals Corporation; Pfizer Inc.; Piramal Imaging; Servier; Takeda Pharmaceutical Company; and Transition Therapeutics. The Canadian Institutes of Health Research is providing funds to support ADNI clinical sites in Canada. Private sector contributions are facilitated by the Foundation for the National Institutes of Health (www.fnih.org). The grantee organization is the Northern California Institute for Research and Education, and the study is coordinated by the Alzheimer’s Therapeutic Research Institute at the University of Southern California. ADNI data are disseminated by the Laboratory for Neuro Imaging at the University of Southern California.

Data used in the preparation of this article were also obtained in part from the Harvard Aging Brain Study (HABS - P01AG036694; https://habs.mgh.harvard.edu). The HABS study was launched in 2010, funded by the National Institute on Aging. and is led by principal investigators Reisa A. Sperling MD and Keith A. Johnson MD at Massachusetts General Hospital/Harvard Medical School in Boston, MA.

OASIS data were provided by OASIS 3: Longitudinal Multimodal Neuroimaging: Principal Investigators: T. Benzinger, D. Marcus, J. Morris; NIH P30 AG066444, P50 AG00561, P30 NS09857781, P01 AG026276, P01 AG003991, R01 AG043434, UL1 TR000448, R01 EB009352. AV-45 doses were provided by Avid Radiopharmaceuticals, a wholly owned subsidiary of Eli Lilly and OASIS-I (test-retest reliability dataset): Cross-Sectional: Principal Investigators: D. Marcus, R, Buckner, J, Csernansky J. Morris; P50 AG05681, P01 AG03991, P01 AG026276, R01 AG021910, P20 MH071616, U24 RR021382. PREVENT-AD was funded by the Canadian Institutes of Health Research, McGill University, the Fonds de Recherche du Québec – Santé, Alzheimer’s Association, Brain Canada, the Government of Canada, the Canada Fund for Innovation, the Douglas Hospital Research Centre and Foundation, the Levesque Foundation, an unrestricted research grant from Pfizer Canada. Private sector contributions are facilitated by the Development Office of the McGill University Faculty of Medicine and by the Douglas Hospital Research Centre Foundation (http://www.douglas.qc.ca/).

UK Biobank is generously supported by its founding funders the Wellcome Trust and UK Medical Research Council, as well as the Department of Health, Scottish Government, the Northwest Regional Development Agency, British Heart Foundation and Cancer Research UK. The organisation has over 150 dedicated members of staff, based in multiple locations across the UK.

Parts of the data are from VETSA, which is funded by the National Institute of Aging grants R01s AG018384, AG018386, AG050595, AG022381, AG076838. The content is the responsibility of the authors and does not necessarily represent official views of the NIA, NIH, or VA. U.S. Department of Veterans Affairs, Department of Defense; National Personnel Records Center, National Archives and Records Administration; Internal Revenue Service; National Opinion Research Center; National Research Council, National Academy of Sciences; and the Institute for Survey Research, Temple University provided invaluable assistance in the conduct of the VET Registry. The Cooperative Studies Program of the U.S. Department of Veterans Affairs provided financial support for development and maintenance of the Vietnam Era Twin Registry. We would also like to acknowledge the continued cooperation and participation of the members of the VET Registry and their families. A list of VETSA investigators and instructions for data access requests are available at https://psychiatry.ucsd.edu/research/programs-centers/vetsa/researchers.html.

## Data and Code Availability

The raw data were gathered from many different datasets. Different agreements are required for each dataset. Most datasets are openly available with prespecified data usage agreements. For some datasets, such as UKB, fees may apply. Requests for LCBC, UB, and COGNORM should be submitted to the corresponding principal investigator. The code to fit the model and generate the figures is available at https://github.com/EdvardGrodem/brain-trajectories.

## Author Contributions

Conceptualization: AF, DP and EG. Data curation: DP. Formal analysis: EG, DP and ØS. Funding acquisition: AF and AB. Investigation: DB-F, AMB, GC, SD, RH, SK, UL, LN, AP-L, CS-P, JS, LW and KW. Methodology: EG, AD, DP, SS, BM and PG. Supervision: BM, AF and AB. Project administration: AF and AB. Resources: AF and AB. Visualization: EG. Writing – original draft: EG, AF and DP. Writing – review and editing: All authors

## A Measuring Linear Change

In this study, we used the correlation between a change estimate and the latent, possibly true, change in a linear time-dependent stochastic dynamical system to quantify sensitivity to detect individualized change over time. Due to the time-varying nature of the dynamical system, these equations are difficult to interpret. Here, we include an alternative description using a linear system. We see that the conclusion from the main paper holds; namely, that the total time span of observations is more important than the number of observations. Additionally, we show that, conditional on the signal-to-noise ratio (SNR), these statements are measurement-modality agnostic. For instance, it does not matter whether the measurements come from a T1-weighted or T2-weighted image, or whether the measurements are of volumes, areas, or thicknesses. We show that the only thing that matters is the signal to noise ratio (SNR) of the expected variance of the change to the measurement noise. These results are, of course, not new and are only restated here for clarity.

Given a linear model, let the observations at times *t*_1_, …, *t*_*n*_ follow

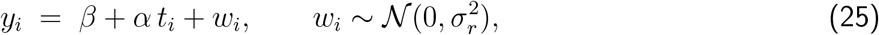

where *α* is the true slope, *β* the intercept, and *σ*_*r*_ the measurement-noise standard deviation (SD). The linear regression slope estimator 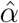 satisfies (Theorem 13.8 of Wasserman, 2004)

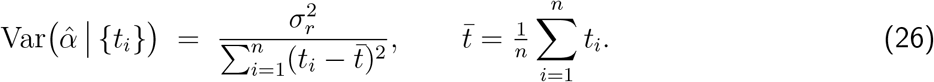

We can see from this that the expected variance of an estimate is inversely proportional to the second central moment of the sample times points.

Given equally spaced (ES) samples, the observations are

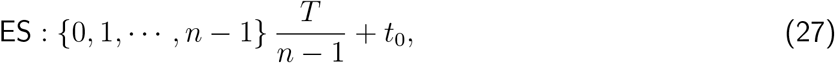

where T is the total span over which the samples are taken. An alternative sampling strategy is to use cluster sampling, where all the samples are at the two endpoints. This assumes, of course, that it is possible to draw independent samples from the same time point. Using cluster sampling (CS) the samples are given by

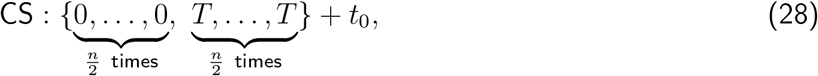

where *n* is assumed to be an even number.

The standard deviation (SD) of the estimator around the true slope for equally spaced samples is then

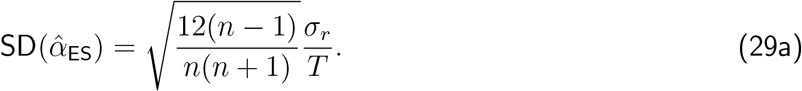

Similarly, for cluster sampling, the variance is

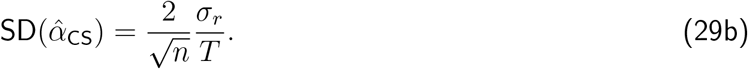

The SNR is the ratio of the variation in slope to the variance of the slope estimate, given by

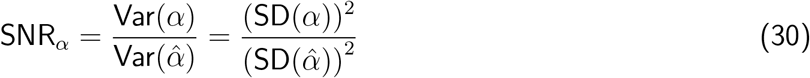

Equation 29 and 30 are visualized in Fig. 7. There are several interesting conclusions that can be drawn from these equations and figures. First, the SD of the slope estimate is proportional to the measurement SD, meaning that halving the measurement SD will halve the slope estimate SD. Second, the SD of the estimate is inversely proportional to the total sampling time, meaning that doubling the total sampling time will halve the SD of the estimate. Both of these conclusions hold for equally spaced sampling and cluster sampling.

**Figure 7:**
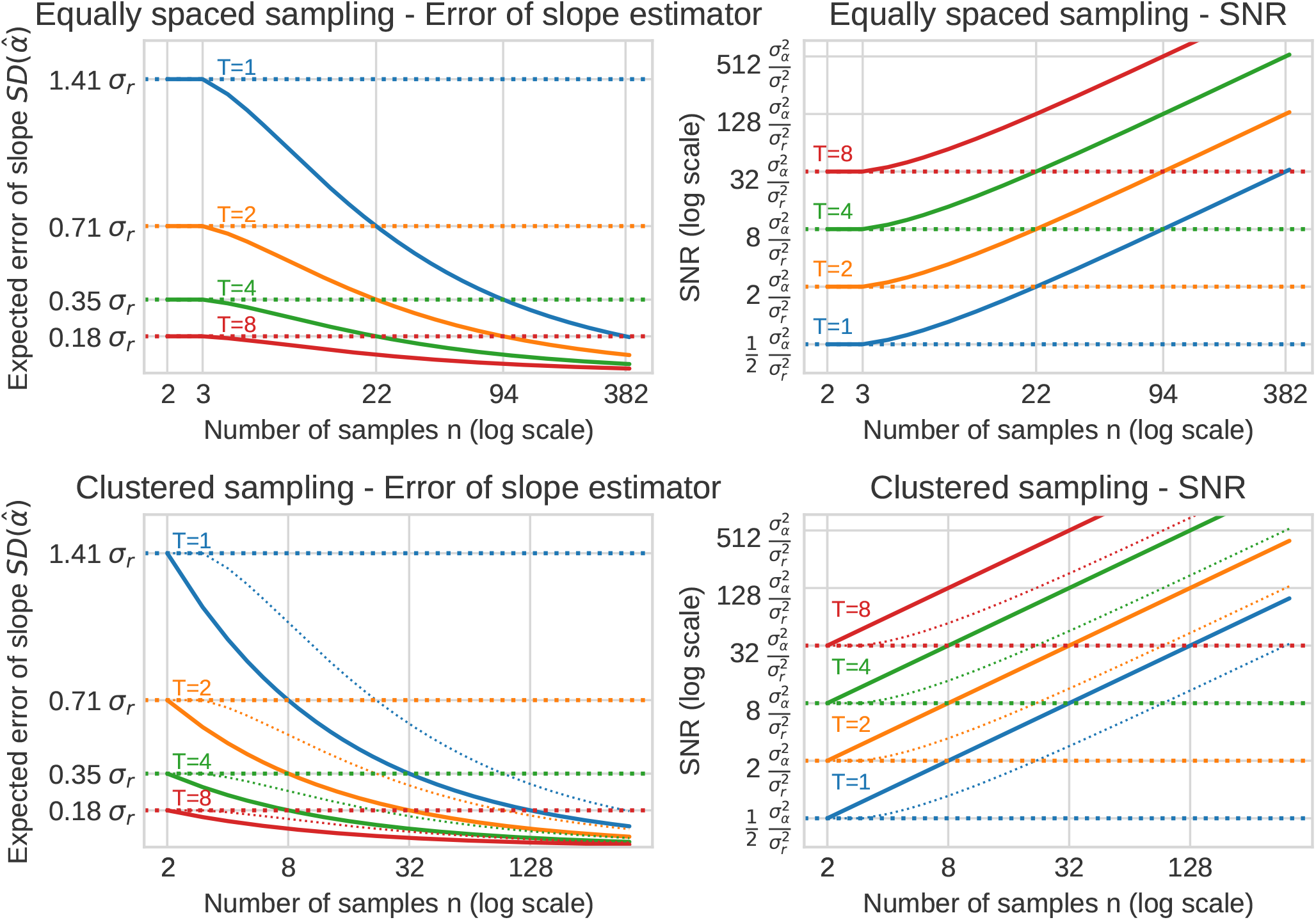
The uncertainty of slope estimate of change with respect to measurement noise, number of samples and the total time span for a linear model of change. To the left the standard deviation (SD) of the slope (SD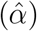) is shown as function of the number of sessions and the total span of the sessions. To the right the signal to noise ratio (SNR) is shown. The horizontal lines show SD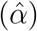 and SNR when *n* = 2. The labels on the x-axis show the number of samples needed for the curves to cross the lines for *n* = 2 of the other values of *T*. In the plots for cluster sampling the thin dotted lines show the curves for equally spaced sampling.

For cluster sampling, the SD of the estimate is inversely proportional to 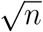, while the story is a little more complicated for equally spaced samples. However, for large n, we have that Var 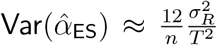, while we know that Var 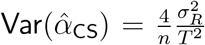, meaning cluster sampling is approximately 3-times more sample efficient for large number of samples. For a smaller number of samples, we can calculate how many samples we need in order to achieve the same improvements as doubling the total sampling time. For cluster sampling we have to increase the number of samples from 2 to 8 samples (4 at each endpoint) in order to achieve the same result as doubling the total sample time. For equally spaced samples, we have to take 22 samples to achieve the same result.

In conclusion, given a linear model with gaussian noise, increasing the total sampling time is more efficient than increasing the number of samples. Cluster sampling is approximately 3 times more sample efficient than equally spaced samples.

## References

Abé, C., Ching, C. R., Liberg, B., Lebedev, A. V., Agartz, I., Akudjedu, T. N., Alda, M., Alnæs, D., Alonso-Lana, S., Benedetti, F., et al. (2022). Longitudinal structural brain changes in bipolar disorder: A multicenter neuroimaging study of 1232 individuals by the ENIGMA bipolar disorder working group. Biological Psychiatry, 91 (6), 582–592.

Abellaneda-Pérez, K., Vaqué-Alcázar, L., Vidal-Piñeiro, D., Jannati, A., Solana, E., Bargalló, N., Santarnecchi, E., Pascual-Leone, A., & Bartrés-Faz, D. (2019). Age-related differences in default-mode network connectivity in response to intermittent theta-burst stimulation and its relationships with maintained cognition and brain integrity in healthy aging. NeuroImage, 188, 794–806.

Allen, J. S., Bruss, J., Brown, C. K., & Damasio, H. (2005). Normal neuroanatomical variation due to age: The major lobes and a parcellation of the temporal region. Neurobiology of Aging, 26 (9), 1245–1260.

Ballester, P. L., Suh, J. S., Ho, N. C., Liang, L., Hassel, S., Strother, S. C., Arnott, S. R., Minuzzi, L., Sassi, R. B., Lam, R. W., et al. (2023). Gray matter volume drives the brain age gap in schizophrenia: A SHAP study. Schizophrenia, 9 (1), 3.

Barnes, J., Ridgway, G. R., Bartlett, J., Henley, S. M., Lehmann, M., Hobbs, N., Clarkson, M. J., MacManus, D. G., Ourselin, S., & Fox, N. C. (2010). Head size, age and gender adjustment in mri studies: A necessary nuisance? NeuroImage, 53 (4), 1244–1255.

Bashyam, V. M., Erus, G., Doshi, J., Habes, M., Nasrallah, I. M., Truelove-Hill, M., Srinivasan, D., Mamourian, L., Pomponio, R., Fan, Y., et al. (2020). MRI signatures of brain age and disease over the lifespan based on a deep brain network and 14 468 individuals worldwide. Brain, 143 (7), 2312–2324.

Battaglini, M., Gentile, G., Luchetti, L., Giorgio, A., Vrenken, H., Barkhof, F., Cover, K. S., Bakshi, R., Chu, R., Sormani, M. P., et al. (2019). Lifespan normative data on rates of brain volume changes. Neurobiology of Aging, 81, 30–37.

Beer, J. C., Tustison, N. J., Cook, P. A., Davatzikos, C., Sheline, Y. I., Shinohara, R. T., Linn, K. A., Initiative, A. D. N., et al. (2020). Longitudinal ComBat: A method for harmonizing longitudinal multi-scanner imaging data. NeuroImage, 220, 117129.

Bertram, L., Böckenhoff, A., Demuth, I., Düzel, S., Eckardt, R., Li, S.-C., Lindenberger, U., Pawelec, G., Siedler, T., Wagner, G. G., et al. (2014). Cohort profile: The Berlin aging study II (BASE-II). International journal of epidemiology, 43 (3), 703–712.

Bingham, E., Chen, J. P., Jankowiak, M., Obermeyer, F., Pradhan, N., Karaletsos, T., Singh, R., Szerlip, P., Horsfall, P., & Goodman, N. D. (2019). Pyro: Deep universal probabilistic programming. Journal of Machine Learning Research, 20 (28), 1–6.

Blake, K. V., Ntwatwa, Z., Kaufmann, T., Stein, D. J., Ipser, J. C., & Groenewold, N. A. (2023). Advanced brain ageing in adult psychopathology: A systematic review and meta-analysis of structural MRI studies. Journal of Psychiatric Research, 157, 180–191.

Brandmaier, A. M., Lindenberger, U., & McCormick, E. M. (2024). Optimal two-time point longitudinal models for estimating individual-level change: Asymptotic insights and practical implications. Developmental Cognitive Neuroscience, 70, 101450.

Brandmaier, A. M., Von Oertzen, T., Ghisletta, P., Lindenberger, U., & Hertzog, C. (2018). Precision, reliability, and effect size of slope variance in latent growth curve models: Implications for statistical power analysis. Frontiers in Psychology, 9, 294.

Brandmaier, A. M., Wenger, E., Bodammer, N. C., Kühn, S., Raz, N., & Lindenberger, U. (2018). Assessing reliability in neuroimaging research through intra-class effect decomposition (ICED). eLife, 7, e35718.

Bråthen, A. C. S., Sørensen, Ø., De Lange, A.-M. G., Mowinckel, A. M., Fjell, A. M., & Walhovd, K. B. (2022). Cognitive and hippocampal changes weeks and years after memory training. Scientific Reports, 12 (1), 7877.

Burgmans, S., van Boxtel, M. P., Gronenschild, E., Vuurman, E., Hofman, P., Uylings, H. B., Jolles, J., & Raz, N. (2010). Multiple indicators of age-related differences in cerebral white matter and the modifying effects of hypertension. NeuroImage, 49 (3), 2083–2093.

Cattaneo, G., Bartrés-Faz, D., Morris, T. P., Sánchez, J. S., Macía, D., Tarrero, C., Tormos, J. M., & Pascual-Leone, A. (2018). The Barcelona brain health initiative: A cohort study to define and promote determinants of brain health. Frontiers in Aging Neuroscience, 10, 321.

Cole, J. H., Ritchie, S. J., Bastin, M. E., Hernández, V., Muñoz Maniega, S., Royle, N., Corley, J., Pattie, A., Harris, S. E., Zhang, Q., et al. (2018). Brain age predicts mortality. Molecular Psychiatry, 23 (5), 1385–1392.

Constantinides, C., Han, L. K., Alloza, C., Antonucci, L. A., Arango, C., Ayesa-Arriola, R., Banaj, N., Bertolino, A., Borgwardt, S., Bruggemann, J., et al. (2023). Brain ageing in schizophrenia: Evidence from 26 international cohorts via the ENIGMA schizophrenia consortium. Molecular Psychiatry, 28 (3), 1201–1209.

Cox, S. R., Dickie, D. A., Ritchie, S. J., Karama, S., Pattie, A., Royle, N. A., Corley, J., Aribisala, B. S., Valdés Hernández, M., Muñoz Maniega, S., et al. (2016). Associations between education and brain structure at age 73 years, adjusted for age 11 IQ. Neurology, 87 (17), 1820–1826.

Dagley, A., LaPoint, M., Huijbers, W., Hedden, T., McLaren, D. G., Chatwal, J. P., Papp, K. V., Amariglio, R. E., Blacker, D., Rentz, D. M., et al. (2017). Harvard aging brain study: Dataset and accessibility. NeuroImage, 144, 255–258.

Dahdah, S., & Forbes, J. R. (2025). Discretization of linear systems using the matrix exponential. arXiv preprint 2505.18187.

DeCarli, C., Maillard, P., Pase, M. P., Beiser, A. S., Kojis, D., Satizabal, C. L., Himali, J. J., Aparicio, H. J., Fletcher, E., & Seshadri, S. (2024). Trends in intracranial and cerebral volumes of Framingham heart study participants born 1930 to 1970. JAMA neurology, 81 (5), 471–480.

Di Biase, M. A., Tian, Y. E., Bethlehem, R. A., Seidlitz, J., Alexander-Bloch, A. F., Yeo, B. T., & Zalesky, A. (2023). Mapping human brain charts cross-sectionally and longitudinally. Proceedings of the National Academy of Sciences, 120 (20), e2216798120.

Dörfel, R. P., Ozenne, B., Ganz, M., Svensson, J., & Plavén-Sigray, P. (2023). Prediction of brain age using structural magnetic resonance imaging: A comparison of accuracy and test–retest reliability of publicly available software packages. Human Brain Mapping, 44 (17), 6139–6148. 10.1002/hbm.26502

Driscoll, I., Davatzikos, C., An, Y., Wu, X., Shen, D., Kraut, M., & Resnick, S. (2009). Longitudinal pattern of regional brain volume change differentiates normal aging from MCI. Neurology, 72 (22), 1906–1913.

Driver, C. C., & Voelkle, M. C. (2018). Hierarchical Bayesian continuous time dynamic modeling. Psychological Methods, 23 (4), 774.

Dugas, C., Bengio, Y., Bélisle, F., Nadeau, C., & Garcia, R. (2000). Incorporating second-order functional knowledge for better option pricing. Advances in Neural Information Processing Systems, 13.

Einstein, A. (1906). On the theory of the Brownian movement. Ann. Phys, 19 (4), 371–381.

Elliott, M. L., Nielsen, J. A., Hanford, L. C., Hamadeh, A., Hilbert, T., Kober, T., Dickerson, B. C., Hyman, B. T., Mair, R. W., Eldaief, M. C., et al. (2024). Precision brain morphometry using cluster scanning. Imaging Neuroscience, 2, 1–15.

Engvig, A., Fjell, A. M., Westlye, L. T., Moberget, T., Sundseth, Ø., Larsen, V. A., & Walhovd, K. B. (2010). Effects of memory training on cortical thickness in the elderly. NeuroImage, 52 (4), 1667– 1676.

Fischl, B., Salat, D. H., Busa, E., Albert, M., Dieterich, M., Haselgrove, C., Van Der Kouwe, A., Killiany, R., Kennedy, D., Klaveness, S., et al. (2002). Whole brain segmentation: Automated labeling of neuroanatomical structures in the human brain. Neuron, 33 (3), 341–355.

Fjell, A. M., Rogeberg, O., Sørensen, Ø., Amlien, I. K., Bartrés-Faz, D., Brandmaier, A. M., Cattaneo, G., Düzel, S., Grydeland, H., Henson, R. N., Kühn, S., Lindenberger, U., Lyngstad, T. H., Mowinckel, A. M., Nyberg, L., Pascual-Leone, A., Solé-Padullés, C., Sneve, M. H., Solana, J., … Vidal-Piñeiro, D. (2025). Reevaluating the role of education on cognitive decline and brain aging in longitudinal cohorts across 33 Western countries. Nature Medicine, 1–10. 10.1038/s41591-025-03828-y

Fjell, A. M., Walhovd, K. B., Fennema-Notestine, C., McEvoy, L. K., Hagler, D. J., Holland, D., Brewer, J. B., & Dale, A. M. (2009). One-year brain atrophy evident in healthy aging. Journal of Neuroscience, 29 (48), 15223–15231.

Fjell, A. M., Westlye, L. T., Grydeland, H., Amlien, I., Espeseth, T., Reinvang, I., Raz, N., Holland, D., Dale, A. M., Walhovd, K. B., et al. (2013). Critical ages in the life course of the adult brain: Nonlinear subcortical aging. Neurobiology of Aging, 34 (10), 2239–2247.

Föllmer, H., & Schied, A. (2011). Stochastic finance: An introduction in discrete time. Walter de Gruyter.

Franke, K., & Gaser, C. (2019). Ten years of BrainAGE as a neuroimaging biomarker of brain aging: What insights have we gained? Frontiers in Neurology, 10, 789.

Friston, K. (2010). The free-energy principle: A unified brain theory? Nature Reviews Neuroscience, 11 (2), 127–138.

Fujita, S., Mori, S., Onda, K., Hanaoka, S., Nomura, Y., Nakao, T., Yoshikawa, T., Takao, H., Hayashi, N., & Abe, O. (2023). Characterization of brain volume changes in aging individuals with normal cognition using serial magnetic resonance imaging. JAMA Network Open, 6 (6), e2318153– e2318153.

Fürtjes, A. E., Foote, I. F., Xia, C., Davies, G., Moodie, J., Taylor, A., Liewald, D. C., Redmond, P., Corley, J., McIntosh, A. M., et al. (2025). Measurement characteristics and genome-wide correlates of lifetime brain atrophy estimated from a single MRI. Nature Communications, 16 (1), 6725.

Gardiner, C. W., et al. (1985). Handbook of stochastic methods (Vol. 3). springer Berlin.

Geyer, C. J. (2011). Introduction to Markov chain Monte Carlo. Handbook of Markov chain Monte Carlo, 20116022 (45), 22.

Ghisletta, P., Mason, F., von Oertzen, T., Hertzog, C., Nilsson, L.-G., & Lindenberger, U. (2020). On the use of growth models to study normal cognitive aging. International Journal of Behavioral Development, 44 (1), 88–96.

Habes, M., Janowitz, D., Erus, G., Toledo, J., Resnick, S., Doshi, J., Van der Auwera, S., Wittfeld, K., Hegenscheid, K., Hosten, N., et al. (2016). Advanced brain aging: Relationship with epidemiologic and genetic risk factors, and overlap with Alzheimer disease atrophy patterns. Translational Psychiatry, 6 (4), e775–e775.

Han, L. K., Dinga, R., Hahn, T., Ching, C. R., Eyler, L. T., Aftanas, L., Aghajani, M., Aleman, A., Baune, B. T., Berger, K., et al. (2021). Brain aging in major depressive disorder: Results from the ENIGMA major depressive disorder working group. Molecular Psychiatry, 26 (9), 5124–5139.

Han, X., Jovicich, J., Salat, D., van der Kouwe, A., Quinn, B., Czanner, S., Busa, E., Pacheco, J., Albert, M., Killiany, R., et al. (2006). Reliability of MRI-derived measurements of human cerebral cortical thickness: The effects of field strength, scanner upgrade and manufacturer. NeuroImage, 32 (1), 180–194.

Hertzog, C. (1985). An individual differences perspective: Implications for cognitive research in gerontology. Research on Aging, 7 (1), 7–45.

Hertzog, C., von Oertzen, T., Ghisletta, P., & Lindenberger, U. (2008). Evaluating the power of latent growth curve models to detect individual differences in change. Structural Equation Modeling: A Multidisciplinary Journal, 15 (4), 541–563.

Hoffman, M. D., Gelman, A., et al. (2014). The No-U-Turn sampler: Adaptively setting path lengths in Hamiltonian Monte Carlo. Journal of Machine Learning Research, 15 (1), 1593–1623.

Hoogman, M., Bralten, J., Hibar, D. P., Mennes, M., Zwiers, M. P., Schweren, L. S., van Hulzen, K. J., Medland, S. E., Shumskaya, E., Jahanshad, N., et al. (2017). Subcortical brain volume differences in participants with attention deficit hyperactivity disorder in children and adults: A cross-sectional mega-analysis. The Lancet Psychiatry, 4 (4), 310–319.

Idland, A.-V., Sala-Llonch, R., Borza, T., Watne, L. O., Wyller, T. B., Brækhus, A., Zetterberg, H., Blennow, K., Walhovd, K. B., & Fjell, A. M. (2017). CSF neurofilament light levels predict hippocampal atrophy in cognitively healthy older adults. Neurobiology of Aging, 49, 138–144.

Kalman, R. E. (1960). A new approach to linear filtering and prediction problems. Journal of Basic Engineering, 82 (1), 35–45. 10.1115/1.3662552

Kennedy, K. M., & Raz, N. (2009). Aging white matter and cognition: Differential effects of regional variations in diffusion properties on memory, executive functions, and speed. Neuropsychologia, 47 (3), 916–927.

Kim, Y. S., Park, I. S., Kim, H. J., Kim, D., Lee, N. J., & Rhyu, I. J. (2018). Changes in intracranial volume and cranial shape in modern koreans over four decades. American Journal of Physical Anthropology, 166 (3), 753–759.

Korbmacher, M., Vidal-Pineiro, D., Wang, M.-Y., van der Meer, D., Wolfers, T., Nakua, H., Eikefjord, E., Andreassen, O. A., Westlye, L. T., & Maximov, I. I. (2025). Cross-sectional brain age assessments are limited in predicting future brain change. Human Brain Mapping, 46 (6), e70203.

Kremen, W. S., Franz, C. E., & Lyons, M. J. (2019). Current status of the vietnam era twin study of aging (VETSA). Twin Research and Human Genetics, 22 (6), 783–787.

LaMontagne, P. J., Benzinger, T. L., Morris, J. C., Keefe, S., Hornbeck, R., Xiong, C., Grant, E., Hassenstab, J., Moulder, K., Vlassenko, A. G., et al. (2019). OASIS-3: Longitudinal neuroimaging, clinical, and cognitive dataset for normal aging and Alzheimer disease. medRxiv, 2019–12.

Lee, U. (1992). Do stock prices follow random walk?: Some international evidence. International Review of Economics & Finance, 1 (4), 315–327.

Lindenberger, U., Von Oertzen, T., Ghisletta, P., & Hertzog, C. (2011). Cross-sectional age variance extraction: What’s change got to do with it? Psychology and Aging, 26 (1), 34.

Livingston, G., Huntley, J., Liu, K. Y., Costafreda, S. G., Selbæk, G., Alladi, S., Ames, D., Banerjee, S., Burns, A., Brayne, C., et al. (2024). Dementia prevention, intervention, and care: 2024 report of the Lancet standing Commission. The Lancet, 404 (10452), 572–628.

Lövdén, M., Garzón, B., & Lindenberger, U. (2020). Human skill learning: Expansion, exploration, selection, and refinement. Current Opinion in Behavioral Sciences, 36, 163–168.

Miller, K. L., Alfaro-Almagro, F., Bangerter, N. K., Thomas, D. L., Yacoub, E., Xu, J., Bartsch, A. J., Jbabdi, S., Sotiropoulos, S. N., Andersson, J. L., et al. (2016). Multimodal population brain imaging in the UK Biobank prospective epidemiological study. Nature Neuroscience, 19 (11), 1523–1536.

Mueller, S. G., Weiner, M. W., Thal, L. J., Petersen, R. C., Jack, C., Jagust, W., Trojanowski, J. Q., Toga, A. W., & Beckett, L. (2005). The Alzheimer’s disease neuroimaging initiative. Neuroimaging Clinics, 15 (4), 869–877.

Murphy, K. P. (2023). Probabilistic machine learning: Advanced topics. MIT Press. http://probml.github.io/book2

Narayanan, S., Nakamura, K., Fonov, V. S., Maranzano, J., Caramanos, Z., Giacomini, P. S., Collins, D. L., & Arnold, D. L. (2020). Brain volume loss in individuals over time: Source of variance and limits of detectability. NeuroImage, 214, 116737.

Nilsson, L.-G., Adolfsson, R., Bäckman, L., de Frias, C. M., Molander, B., & Nyberg, L. (2004). Betula: A prospective cohort study on memory, health and aging. Aging Neuropsychology and Cognition, 11 (2-3), 134–148.

Nobis, L., Manohar, S. G., Smith, S. M., Alfaro-Almagro, F., Jenkinson, M., Mackay, C. E., & Husain, M. (2019). Hippocampal volume across age: Nomograms derived from over 19,700 people in UK Biobank. NeuroImage: Clinical, 23, 101904.

Nocedal, J., & Wright, S. J. (2006). Numerical optimization. Sprinter.

Nyberg, L., Boraxbekk, C.-J., Sörman, D. E., Hansson, P., Herlitz, A., Kauppi, K., Ljungberg, J. K., Lövheim, H., Lundquist, A., Adolfsson, A. N., et al. (2020). Biological and environmental predictors of heterogeneity in neurocognitive ageing: Evidence from Betula and other longitudinal studies. Ageing Research Reviews, 64, 101184.

Nyberg, L., Magnussen, F., Lundquist, A., Baaré, W., Bartrés-Faz, D., Bertram, L., Boraxbekk, C.-J., Brandmaier, A. M., Drevon, C. A., Ebmeier, K., et al. (2021). Educational attainment does not influence brain aging. Proceedings of the National Academy of Sciences, 118 (18), e2101644118.

Nyberg, L., Salami, A., Andersson, M., Eriksson, J., Kalpouzos, G., Kauppi, K., Lind, J., Pudas, S., Persson, J., & Nilsson, L.-G. (2010). Longitudinal evidence for diminished frontal cortex function in aging. Proceedings of the National Academy of Sciences, 107 (52), 22682–22686.

Pfefferbaum, A., Rohlfing, T., Rosenbloom, M. J., Chu, W., Colrain, I. M., & Sullivan, E. V. (2013). Variation in longitudinal trajectories of regional brain volumes of healthy men and women (ages 10 to 85 years) measured with atlas-based parcellation of MRI. NeuroImage, 65, 176–193.

Preacher, K. J. (2008). Latent growth curve modeling. Sage.

Rajaram, S., Valls-Pedret, C., Cofán, M., Sabaté, J., Serra-Mir, M., Pérez-Heras, A. M., Arechiga, A., Casaroli-Marano, R. P., Alforja, S., Sala-Vila, A., et al. (2017). The Walnuts and Healthy Aging Study (WAHA): Protocol for a nutritional intervention trial with walnuts on brain aging. Frontiers in Aging Neuroscience, 8, 333.

Rauch, H. E., Tung, F., & Striebel, C. T. (1965). Maximum likelihood estimates of linear dynamic systems. AIAA Journal, 3 (8), 1445–1450.

Raz, N., & Lindenberger, U. (2011). Only time will tell: Cross-sectional studies offer no solution to the age–brain–cognition triangle: Comment on Salthouse (2011). Psychological Bulletin.

Raz, N., Lindenberger, U., Rodrigue, K. M., Kennedy, K. M., Head, D., Williamson, A., Dahle, C., Gerstorf, D., & Acker, J. D. (2005). Regional brain changes in aging healthy adults: General trends, individual differences and modifiers. Cerebral Cortex, 15 (11), 1676–1689.

Reuter, M., Schmansky, N. J., Rosas, H. D., & Fischl, B. (2012). Within-subject template estimation for unbiased longitudinal image analysis. NeuroImage, 61 (4), 1402–1418.

Rohrer, J. M. (2025). Thinking clearly about age, period, and cohort effects. Advances in Methods and Practices in Psychological Science, 8 (2), 25152459251342750.

Schwarz, C. G., Gunter, J. L., Wiste, H. J., Przybelski, S. A., Weigand, S. D., Ward, C. P., Senjem, M. L., Vemuri, P., Murray, M. E., Dickson, D. W., et al. (2016). A large-scale comparison of cortical thickness and volume methods for measuring Alzheimer’s disease severity. NeuroImage: Clinical, 11, 802–812.

Sele, S., Liem, F., Mérillat, S., & Jäncke, L. (2021). Age-related decline in the brain: A longitudinal study on inter-individual variability of cortical thickness, area, volume, and cognition. NeuroImage, 240, 118370.

Shafto, M. A., Tyler, L. K., Dixon, M., Taylor, J. R., Rowe, J. B., Cusack, R., Calder, A. J., Marslen-Wilson, W. D., Duncan, J., Dalgleish, T., et al. (2014). The Cambridge Centre for Ageing and Neuroscience (Cam-CAN) study protocol: A cross-sectional, lifespan, multidisciplinary examination of healthy cognitive ageing. BMC neurology, 14, 1–25.

Simon, D. (2006). Optimal state estimation: Kalman, H infinity, and nonlinear approaches. John Wiley & Sons.

Smith, S. M., Elliott, L. T., Alfaro-Almagro, F., McCarthy, P., Nichols, T. E., Douaud, G., & Miller, K. L. (2020). Brain aging comprises many modes of structural and functional change with distinct genetic and biophysical associations. eLife, 9, e52677.

Smith, S. M., Miller, K. L., & Nichols, T. E. (2025). Characterising ongoing brain aging and baseline effects from cross-sectional data. Imaging Neuroscience, 3, IMAG.a.39. 10.1162/IMAG.a.39

Smith, S. M., Vidaurre, D., Alfaro-Almagro, F., Nichols, T. E., & Miller, K. L. (2019). Estimation of brain age delta from brain imaging. NeuroImage, 200, 528–539.

Sørensen, Ø., & McCormick, E. M. (2025). Modeling cycles, trends and time-varying effects in dynamic structural equation models with regression splines. Multivariate Behavioral Research, 60 (5), 1013– 1028.

Tremblay-Mercier, J., Madjar, C., Das, S., Binette, A. P., Dyke, S. O., Étienne, P., Lafaille-Magnan, M.-E., Remz, J., Bellec, P., Collins, D. L., et al. (2021). Open science datasets from PREVENTAD, a longitudinal cohort of pre-symptomatic Alzheimer’s disease. NeuroImage: Clinical, 31, 102733.

Uribe, C., Segura, B., Baggio, H. C., Abos, A., Marti, M. J., Valldeoriola, F., Compta, Y., Bargallo, N., & Junque, C. (2016). Patterns of cortical thinning in nondemented Parkinson’s disease patients. Movement Disorders, 31 (5), 699–708.

Van Erp, T. G., Walton, E., Hibar, D. P., Schmaal, L., Jiang, W., Glahn, D. C., Pearlson, G. D., Yao, N., Fukunaga, M., Hashimoto, R., et al. (2018). Cortical brain abnormalities in 4474 individuals with schizophrenia and 5098 control subjects via the enhancing neuro imaging genetics through meta analysis (ENIGMA) consortium. Biological Psychiatry, 84 (9), 644–654.

Vidal-Pineiro, D., Wang, Y., Krogsrud, S. K., Amlien, I. K., Baaré, W. F., Bartres-Faz, D., Bertram, L., Brandmaier, A. M., Drevon, C. A., Düzel, S., et al. (2021). Individual variations in ‘brain age’ relate to early-life factors more than to longitudinal brain change. eLife, 10, e69995.

Vidal-Piñeiro, D., Sørensen, Ø., Strømstad, M., Amlien, I. K., Anderson, M., Baaré, W. F., Bartrés-Faz, D., Brandmaier, A. M., Bråthen, A. C., Garrido, P., et al. (2025). Reliability of structural brain change in cognitively healthy adult samples. Imaging Neuroscience.

Vidal-Piñeiro, D., Martin-Trias, P., Arenaza-Urquijo, E. M., Sala-Llonch, R., Clemente, I. C., Mena-Sánchez, I., Bargalló, N., Falcón, C., Pascual-Leone, Á., & Bartrés-Faz, D. (2014). Task-dependent activity and connectivity predict episodic memory network-based responses to brain stimulation in healthy aging. Brain Stimulation, 7 (2), 287–296.

Vita, A., De Peri, L., Deste, G., & Sacchetti, E. (2012). Progressive loss of cortical gray matter in schizophrenia: A meta-analysis and meta-regression of longitudinal MRI studies. Translational Psychiatry, 2 (11), e190–e190.

Walhovd, K. B., Fjell, A. M., Brown, T. T., Kuperman, J. M., Chung, Y., Hagler Jr, D. J., Roddey, J. C., Erhart, M., McCabe, C., Akshoomoff, N., et al. (2012). Long-term influence of normal variation in neonatal characteristics on human brain development. Proceedings of the National Academy of Sciences, 109 (49), 20089–20094.

Walhovd, K. B., Fjell, A. M., Reinvang, I., Lundervold, A., Dale, A. M., Eilertsen, D. E., Quinn, B. T., Salat, D., Makris, N., & Fischl, B. (2005). Effects of age on volumes of cortex, white matter and subcortical structures. Neurobiology of Aging, 26 (9), 1261–1270.

Walhovd, K. B., Fjell, A. M., Westerhausen, R., Nyberg, L., Ebmeier, K. P., Lindenberger, U., Bartrés-Faz, D., Baaré, W. F., Siebner, H. R., Henson, R., et al. (2018). Healthy minds 0–100 years: Optimising the use of European brain imaging cohorts (“Lifebrain”). European Psychiatry, 50, 47–56.

Walhovd, K. B., Krogsrud, S. K., Amlien, I. K., Bartsch, H., Bjørnerud, A., Due-Tønnessen, P., Grydeland, H., Hagler Jr, D. J., Håberg, A. K., Kremen, W. S., et al. (2016). Neurodevelopmental origins of lifespan changes in brain and cognition. Proceedings of the National Academy of Sciences, 113 (33), 9357–9362.

Walhovd, K. B., Lövden, M., & Fjell, A. M. (2023). Timing of lifespan influences on brain and cognition. Trends in Cognitive Sciences, 27 (10), 901–915.

Walhovd, K. B., Nyberg, L., Lindenberger, U., Amlien, I. K., Sørensen, Ø., Wang, Y., Mowinckel, A. M., Kievit, R. A., Ebmeier, K. P., Bartrés-Faz, D., et al. (2022). Brain aging differs with cognitive ability regardless of education. Scientific Reports, 12 (1), 13886.

Walhovd, K. B., Westlye, L. T., Amlien, I., Espeseth, T., Reinvang, I., Raz, N., Agartz, I., Salat, D. H., Greve, D. N., Fischl, B., et al. (2011). Consistent neuroanatomical age-related volume differences across multiple samples. Neurobiology of Aging, 32 (5), 916–932.

Wasserman, L. (2004). All of statistics: A concise course in statistical inference. Springer.

Welch, G., Bishop, G., et al. (1995). An introduction to the Kalman filter.

Wenger, E., Kühn, S., Verrel, J., Mårtensson, J., Bodammer, N. C., Lindenberger, U., & Lövdén, M. (2017). Repeated structural imaging reveals nonlinear progression of experience-dependent volume changes in human motor cortex. Cerebral Cortex, 27 (5), 2911–2925.

Wenger, E., Schaefer, S., Noack, H., Kühn, S., Mårtensson, J., Heinze, H.-J., Düzel, E., Bäckman, L., Lindenberger, U., & Lövdén, M. (2012). Cortical thickness changes following spatial navigation training in adulthood and aging. NeuroImage, 59 (4), 3389–3397.

Zhu, J.-D., Wu, Y.-F., Tsai, S.-J., Lin, C.-P., & Yang, A. C. (2023). Investigating brain aging trajectory deviations in different brain regions of individuals with schizophrenia using multimodal magnetic resonance imaging and brain-age prediction: A multicenter study. Translational Psychiatry, 13 (1), 82.

