## Supplementary Materials for "Stable Individual Differences Dominate Adult Brain Volume Variation Until Later Life"

#### 1 Data

##### 1.1 ADNI

The Alzheimer's Disease Neuroimaging Initiative (ADNI) (Mueller et al., 2005) is a multi-site project led by Doctor Michael W. Weiner to assess the progression of mild cognitive impairment (MCI) and early Alzheimer's Disease (AD), combining imaging, clinical and other biological markers, and neuropsychological and clinical assessments over time. For more information, visit https://adni.loni.usc.edu/about/. The age range for the participants is 55-90 years. The present study includes participants from ADNI 1, ADNIGO, ADNI2, and ADNI 3, who were cognitively healthy at baseline (DX_bl variable). Only observations in which participants were still cognitively healthy were included as determined by the ADNI team (DX variable). Amongst others, participants were required to have no evidence of ischemic stroke (Hachinski Ischemic Score $\leq 4$), a Geriatric Depression scale score $< 6$, stable medications for 4 weeks before the screening, good auditory and visual acuity, good general health, no medical contraindications to MRI and at least 6 grades of education/work history. In-detailed general inclusion and exclusion criteria are described elsewhere (Petersen et al., 2010). All participants signed an informed consent form and the protocols were approved by the corresponding regional ethical committees in the US and Canada.

##### 1.2 BASE-II

BASE-II: Participants of the Berlin Aging Study II were community-dwelling older adults recruited from the greater Berlin metropolitan area through advertisements in newspapers and public areas. The baseline sample comprised 2200 participants; 1600 older adults aged 61–88 years, and 600 younger adults aged

24–40 years. Participants were invited to a medical exam and cognitive testing sessions. After completion of the cognitive examination of BASE-II, eligible participants were invited to take part in one MRI session within a time window of 2–4 weeks after cognitive testing. The MRI sample consisted of 341 older adults aged 61–82 years and 103 younger adults. MR scans and cognitive scores were obtained 2012-2013. A subsample of the MR sample was later invited for follow-up. The different elements of the study were approved by the ethics committees of the Max Planck Institute for Human Development, the Charité University ethics committee and by the ethics committees of The German Association for Psychology (DGPs). Participants signed written informed consent and received monetary compensation for their participation in BASE-II and the MRI study. Exclusion criteria were untreated diabetes and hypertension; prior stroke, head injuries or brain surgery; psychiatric illness; major depression; dementia with a score $< 24$ on the Mini-Mental State Examination. None of the participants took medication that might affect memory function or had a history of head injuries, medical (e.g., heart attack), neurological (e.g., epilepsy), or psychiatric disorders (e.g., depression). All participants reported normal or corrected to normal vision. Observations with MMSE $< 26$ or no MMSE data were discarded.

#### 1.3 BBHI

Barcelona Brain Health Initiative study (https://bbhi.cat/en/) participants are community-dwelling individuals between 40 and 65 years of age, without self-reported neurological or psychiatric diagnosis at the time of recruitment 2. BBHI is an ongoing longitudinal cohort study that investigates the determinants of brain and mental health in healthy middle-aged and older adults. Recruitment started in 2017, when multiple initiatives (including conferences, radio and television interviews, and social media advertisements) took place to encourage participants to join the study. It has enrolled 4,686 participants via a web-based application, who completed a first online questionnaire. Exclusion criteria included cognitive impairment and diagnosis of neurologic or psychiatric disorders, including Alzheimer's disease, Parkinson's disease, multiple sclerosis, amyotrophic lateral sclerosis, cerebral stroke, schizophrenia, and major depression. BBHI includes regular cognitive, medical, brain imaging, and biological assessments, and has several sub studies. A sample of 1,000 participants is undergoing a detailed clinical phenotyping through a multi-day in-person evaluation that includes cognitive, physical, and medical assessments, biological sample recollection, structural and functional magnetic resonance imaging (MRI), and electroencephalography (EEG). Participants of this study are invited biannually for repeated evaluations.

#### 1.4 BETULA

The BETULA project (Nilsson et al., 2004) is a prospective longitudinal study on aging, memory, and dementia, which used a population-based sampling of healthy middle-aged and older adults for recruitment. Detailed recruitment procedures are found elsewhere (Nilsson et al., 2004; Nyberg et al., 2020). For the current analyses, the MRI subsample of the study is used. Participation in the neuroimaging study was offered to all participants who had remained in the study and completed cognitive testing at the 5th Betula test wave onwards. Exclusion criteria were severe visual or auditory handicaps, intellectual or developmental disabilities, suspected dementia, having a mother tongue other than Swedish, MRI contraindications, severe neurological disorders, or visual/motor deficits that could interfere with fMRI data collection, MMSE <24, brain or head surgery, and substantial brain anatomical deviations. Some participants were later excluded due to discovered neurological conditions, severe depression, and MRI anatomical abnormalities. The LCBC cohort was part of the Lifebrain obtained as part of the Lifebrain consortium (Walhovd et al., 2018). All participants signed an informed consent and the protocols were approved by the Regional Ethical Vetting Board at Umeå University.

#### 1.5 Cam-CAN

The Cambridge Centre for Ageing and Neuroscience cohort study Shafto et al., 2014 is a large-scale, multi-modal, population-based adult lifespan (18–87 years old) investigation of the neural underpinnings of successful cognitive ageing. Recruitment was done by invitation letters based on the patient lists of general practitioners within the Cambridge City area. A population-based cohort of 2700 adults aged 18 or above was recruited to Stage 1 of the project, where they completed an interview including health and lifestyle questions, a core cognitive assessment, and a questionnaire of lifetime experiences and physical activity. Approximately 700 participants continued to Stage 2 where they underwent cognitive testing and provided measures of brain structure and function. In stage 3, a subset of approximately 250 adults returned for longitudinal follow-up MRI. The study is conducted in compliance with the Helsinki Declaration, and has been approved by the local ethics committee, Cambridgeshire 2. Exclusion criteria included term-time residents of colleges and universities, and participants whose Primary Care Physician judged as inappropriate to include. For phase-II, exclusion criteria additionally included cognitive impairment (MMSE $<$ 24, memory deficit, consent difficulties), communication difficulties (hearing problems, insufficient English language, vision difficulties), medical problems by self-report of diagnosis (dementia diagnosis /Alzheimer's Disease, Parkinson's Disease, Motor Neuron disease, Multiple sclerosis, cancer, stroke, encephalitis, meningitis, epilepsy, head injury with serious results [coma, unconscious for >2 hours, skull fracture], recently diagnosed or uncontrolled high blood pressure, possible pregnancy, current psychiatric conditions [bipolar disorder, schizophrenia, psychosis]), mobility problems (restricted mobility which could prevent further participation, inability to walk 10 meters), substance abuse (past or current treatment for drug abuse, current drug usage), and MRI/ MEG safety and comfort exclusions.

#### 1.6 COGNORM

The COGNORM cohort Idland et al., 2017 is an ongoing, prospective study coordinated by the Oslo University Hospital and Diakonhjemmet Hospital, Oslo, Norway. Patients (age $\geq$ 65 years) scheduled for elective gynecological, urological, or orthopedic surgery under spinal anesthesia were recruited. Participants were required to have no dementia, previous stroke with sequela, Parkinson's disease, or other neurodegenerative diseases that are likely to affect cognition. Patients with suspected undiagnosed dementia at any time within the first five years of follow-up ($n = 15$) (Sajjad et al., 2020), MMSE score $<$ 28 at baseline, and at least two abnormal cognitive test scores (-1.5 standard deviation [SD] below the mean normal value for age, sex, and education) were excluded. All observations corresponding to cognitively normal individuals were included. All participants signed an informed consent form and the protocol was approved by the Norwegian Regional Committees for Medical and Health Research Ethics and the Data Protector Officer at Oslo University Hospital.

#### 1.7 HABS

The Harvard Aging Brain Study (HABS) (Dagley et al., 2017) is an ongoing, long-term observational study that aims to enhance our understanding of brain aging and the early stages of Alzheimer's disease. The study collects PET, MRI data, neuropsychological and clinical assessments. The age range was between 50 and 90 years at the time of baseline assessment and all patients were considered non-clinically impaired at the start of the study. Further participants had a CDR score of 0, MMSE score $\geq$ 25, $<$ 11 on the Geriatric Depression Scale, and scores above age- and education-adjusted cutoffs on the 30-Minute Delayed Recall of the Logical Memory Story A to be included in the study. Participants with a history of alcoholism, drug abuse, head trauma, or current serious medical/psychiatric illness were excluded. Further details can be found elsewhere (Dagley et al., 2017). Observations with MCI or AD diagnostic (DX variable) were excluded. All participants signed an informed consent form and the protocol was approved by the Partners Healthcare Human Research Committee.

#### 1.8 LCBC

The Center for Lifespan Changes in Brain and Cognition cohort (LCBC, Oslo) (Walhovd et al., 2016) consists of cognitively healthy, community-dwelling participants across the lifespan and is drawn from studies coordinated by the LCBC Research Group (LCBC www.oslobrains.no), approved by a Norwegian Regional Committee for Medical and Health Research Ethics. Written informed consent was obtained from all participants. The samples were recruited by a variety of methods such as newspapers and webpage ads. Most participants were recruited for observational studies, some currently ongoing, while a minority were recruited to enter into cognitive training. Written informed consent was obtained from all adult participants. All participants had to undergo a standardized health interview before being included in the study, and those with a history of neurological or psychiatric conditions or who reported concerns about their cognitive function were excluded. Additionally, all participants over the age of 40 years were required to score at least 25 on the Mini-Mental State Examination. The LCBC cohort was part of the Lifebrain obtained as part of the Lifebrain consortium (Walhovd et al., 2018). MRI observations paired with MMSE $\leq$ 25 were excluded.

#### 1.9 OASIS3

The Open Access Series of Imaging Studies (OASIS3) (LaMontagne et al., 2019) is a retrospective collection of multimodal data that focuses on aging and AD and is openly accessible to the scientific community. OASIS-3 includes neuroimaging, clinical and neuropsychological data. Participants were recruited through the Washington University Knight Alzheimer Disease Research Center via flyers, word of mouth, and community engagements and were aged between 42 and 95 years. Only participants deemed cognitively normal at baseline were included in the observations. Exclusion criteria included medical conditions that precluded longitudinal participation or medical contraindications for the different study arms. See in-detail inclusion and exclusion criteriaLaMontagne et al., 2019. All participants consented to Knight ADRC-related projects following procedures approved by the Institutional Review Board of Washington University School of Medicine. Observations were included until the last observation in which a subject was deemed cognitively healthy as determined by the Clinical Dementia Rating Scale (CDR) (Morris, 1993).

#### 1.10 PreventAD

The Pre-symptomatic Evaluation of Experimental or Novel Treatments for AD (PREVENT-AD) (Tremblay-Mercier et al., 2021) is a retrospective, long-term study that follows cognitively healthy older individuals with a familiar history of AD. It includes participants enrolled either from an observational cohort or the clinical trial of PREVENT-AD. This study comprises MRI images, blood and CSF samples, and clinical and neuropsychological assessments. Participants in the study had to be at least 60 years old, had $\geq$ 6 years of education, and they needed to be cognitively unimpaired at baseline. The Montreal Cognitive Assessment (MoCA) and CDR scales were used to assess cognitive abilities, and participants were considered cognitively intact if their MoCA scores were $\geq 26/30$ or their CDR was $= 0$. Other exclusion criteria at baseline included medical conditions that prevented longitudinal participation or medical contraindications to MRI, use of acetylcholinesterase inhibitors, other approved prescription cognitive enhancers, hypertension, or substance abuse. The inclusion and exclusion criteria have been previously described in detail (Tremblay-Mercier et al., 2021). The protocols, consent forms, and study procedures were approved by the McGill Institutional Review Board and the Douglas Mental Health University Institute Research Ethics Board. Observations with RBANS $>$ 1SD below the mean and probable MCI, as evaluated by a clinician, were excluded.

## 1.11 UB

UB: The University of Barcelona cohort consisted of a series of retrospective sub studies, comprising cognitively healthy, community-dwelling participants with normal visual function in the age range 38-89. Most were recruited for observational studies while a minority were recruited to cognitive training. Exclusion criteria varied across sub-studies, but included severe neurologic and psychiatric disorders, recent head trauma or brain surgery, cognitive deterioration, and a score $<$ 24 on the Mini-Mental State Examination. All participants signed informed consent, and the protocols were approved by the ethical committees of the University of Barcelona and of the Hospital Clinic of Barcelona.

#### 1.12 UKB

The UK Biobank (UKB) (https://www.ukbiobank.ac.uk/about-biobank-uk/) is a major national and international health resource with the aim of improving the prevention, diagnosis and treatment of a wide range of illnesses. UK Biobank recruited $\approx$500,000 people aged between 40-69 years in 2006-2010 from across the country to take part in this project (Guggenheim et al., 2015). Potential participants were identified through National Health Service (NHS) registers according to being aged 40-69 and living within a reasonable traveling distance of an assessment center. Assessment centers are located in accessible and convenient locations with a large surrounding population. Participants have undergone measures and provided samples and detailed information about themselves and agreed to have their health followed. The study sample was drawn from the UK Biobank neuroimaging branch Miller et al., 2016 and conducted under data application number 32048. Only individuals with longitudinal MRI data were used in this study. The subsample used in this study consists of the participants in the first wave of longitudinal imaging. Participants signed an informed consent and the protocols were approved by the North West Multi-Center Research Ethics Committee [MREC]; see also https://www.ukbiobank.ac.uk/the-ethics-and-governance-council.

#### 1.13 VETSA

The Vietnam Era Twin Study of Aging (Kremen et al., 2019) is an ongoing large-scale investigation of cognitive and brain aging in men, investigating genetic and environmental influences on cognitive aging, brain structure and function, and health. VETSA involves over 1600 male twins from the Vietnam Era Twin Registry who served during the Vietnam War era, between 1965 and 1975, though approximately 80% report no combat experience. Assessments began when participants were in their 50s (in 2003) and follow-ups are conducted every 5-6 years. The age range is 52 to 60 at baseline. Assessments include extensive neurocognitive testing, genetics, brain MRI, and plasma samples. $>$ 1200 twins participated in waves 1, 2 and 3. The sample is relatively representative of US men in their age range. Attrition-replacement procedures were taken in wave 2. For the VETSA MRI study, participants are screened for safety issues (e.g. MRI contraindications), and both members of a twin pair had to consent to participate. For MRI, twins had to additionally be able to travel to a scanning site. Other exclusion criteria were depended on exclusion criteria for serving in the military, e.g. participants scoring in the lowest 10 percentile ranks of the Armed Forced Qualification Test (AFQT) were excluded from the military. In addition, we further excluded data from participants with incidental radiological findings, history of seizure, and diagnostic of multiple sclerosis, and AIDS.

#### 1.14 Wayne

The Wayne cohort (Burgmans et al., 2010; Kennedy & Raz, 2009) consists of two retrospective longitudinal datasets which are part of the Brain Aging in Detroit Longitudinal Study. The overarching aim of the study is to understand the mechanisms driving human brain changes over the adult lifespan, identify the risk factors and protective influences that modify the rate of change, and elucidate the relationships between changes in brain properties and cognitive performance. See a detailed description of the sample and inclusion criteria elsewhere (Burgmans et al., 2010; Kennedy & Raz, 2009). Briefly, all participants were screened at baseline via a health questionnaire for the following conditions: the presence of cardiovascular, neurological, or psychiatric disease, use of centrally acting medications, the habit of having three or more alcoholic drinks per day, as well as being minimally high school educated, native English speakers were inclusion/exclusion criteria. Healthy volunteers from the metropolitan Detroit area who responded to media advertisements and flyers participated in this study. Further exclusion criteria included MMSE $<$ 26 and GDS $>$16. All participants provided written informed consent. The study protocols were approved by the (Wayne State) University Institutional Review Board.

### 2 Priors

Before fitting the model each structure volume is z-scored such that the whole dataset population has a mean of 0 and a std of 1. The priors are set in z-scored units such that the same priors relative to the scale of each structure are used. The time is normalized to [0, 1] such that, for instance, the prior of the initial change rate $\beta_{v0} \sim \mathcal{N}(0, 5)$ corresponds to a prior covering the change rates from -10 std to $+$10 std over the lifespan when considering the 95% CI. Notice that the priors on the $b(t)$ correspond to the common L2 smoothing priors for splines. Several of the scale terms used a prior set by a Normal distribution transformed by a Softplus function, defined as Softplus$(x) = ln(e^x + 1)$.

### 3 Verification of Invariance to Dataset Distribution

In Fig. **??** as well as in Fig. **??** one can notice that the age group from 50 to 70 has a higher representation than the younger age group. We also see that the spread of the variance in the change rate occurs at the beginning of the high density data. To ensure that this is not an artifact of the age distribution of the sample, we performed a second fit to a subsample of the data. The subsample consisted of the LCBC dataset and the Cam-CAN dataset, both of which have an even age distribution. We observe that the confidence intervals for the normative curves are wider than those for the full dataset, as expected. However, we see no significant differences in the conclusions drawn from the full dataset. We therefore conclude that the spread of change rate variance at age 50 is not due to an artifact of age distribution.

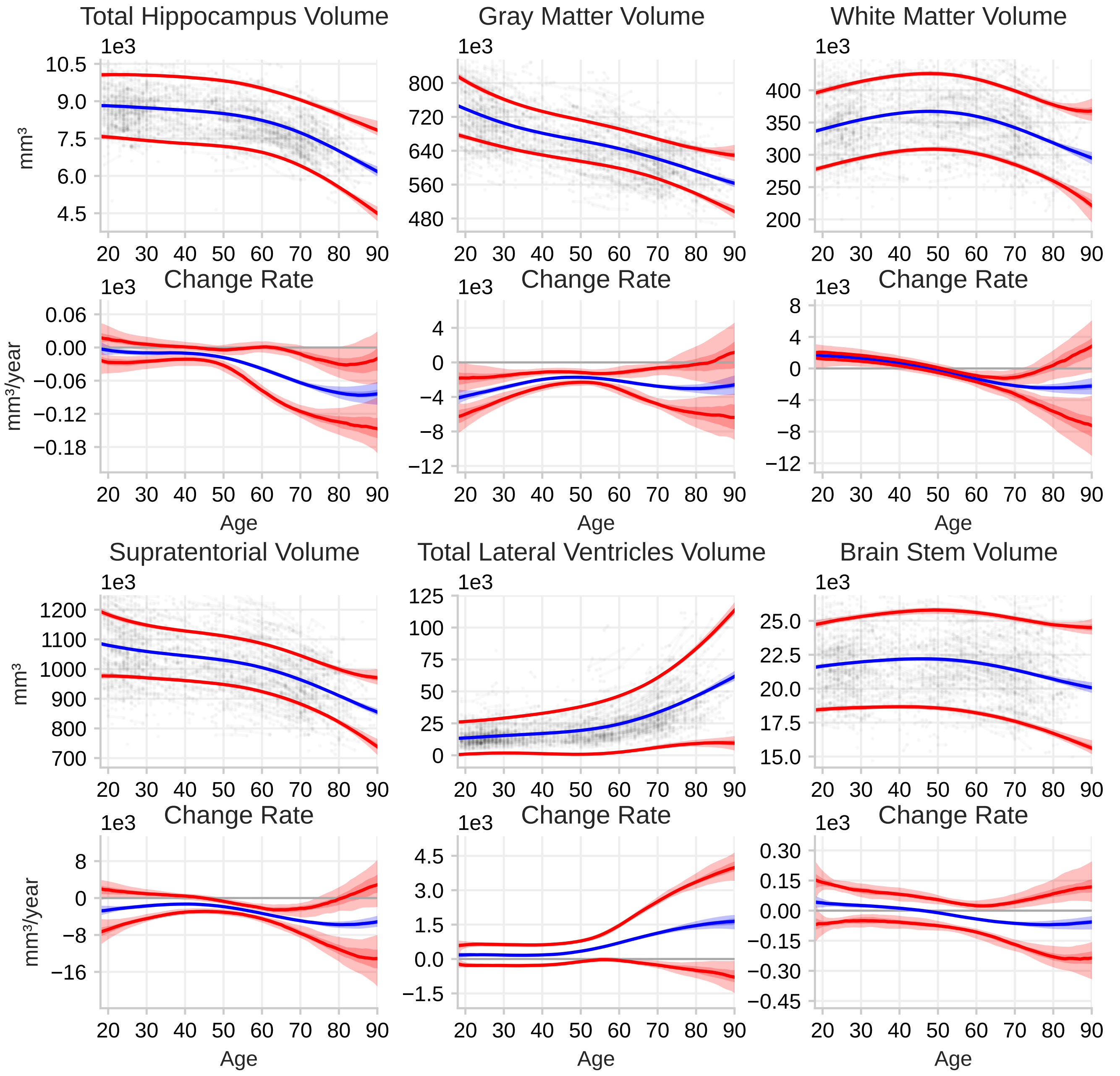


Figure 1: Reproduction of Fig. 3 in the main paper. However here only data from the LCBC dataset and Cam-CAN is used to fit the model such that the age distribution is more uniform. The CIs around the lines are much wider, especially for older ages and the axes are adjusted to include the full CIs.

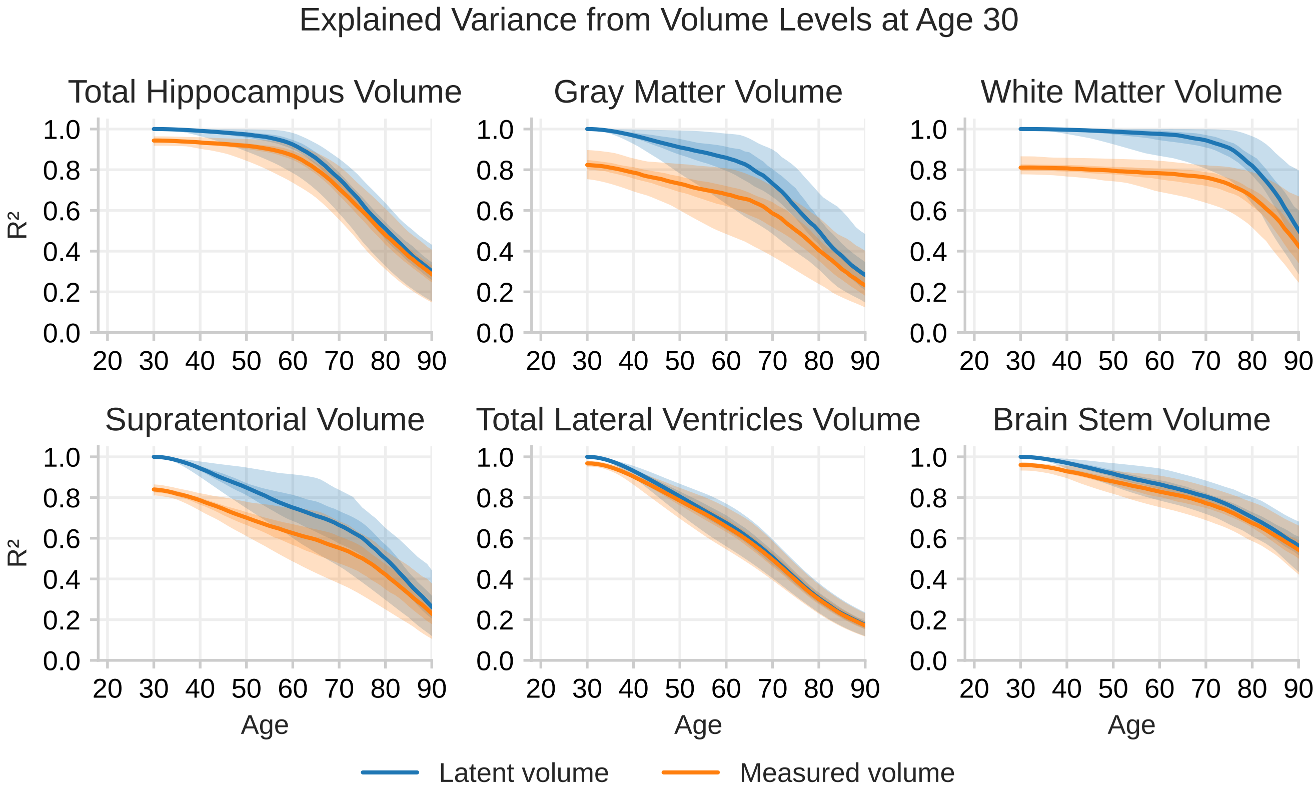


Figure 2: Reproduction of Fig. 4 in the main paper. However here only data from the LCBC dataset and Cam-CAN are used to fit the model.

Table 1: Priors for the model parameters.

| Parameter | Prior Distribution | Comments |
|---|---|---|
| $b(t)$ | $\mathcal{N}(0, 10)$ | |
| $\sigma_q(t)$ | $\text{Softplus}\Big(\mathcal{N}(a, b)\Big)$ | such that $E[\sigma_q] \approx 10$, $Std[\sigma_q] \approx 10$ |
| $\beta_{s0}$ | $\mathcal{N}(0, 5)$ | |
| $\beta_{v0}$ | $\mathcal{N}(0, 5)$ | |
| $\sigma_{s0}$ | $\text{Softplus}\Big(\mathcal{N}(a, b)\Big)$ | such that $E[\sigma_{s0}] \approx 2$, $Std[\sigma_{s0}] \approx 2$ |
| $\sigma_{v0}$ | $\text{Softplus}\Big(\mathcal{N}(a, b)\Big)$ | such that $E[\sigma_{v0}] \approx 2$, $Std[\sigma_{v0}] \approx 2$ |
| $\beta_{s0,\text{sex}}$ | $\mathcal{N}(0, 0.2)$ | |
| $\beta_{s0,\text{ICV}}$ | $\mathcal{N}(0, 1.0)$ | |
| $\beta_{v0,\text{sex}}$ | $\mathcal{N}(0, 0.01)$ | |
| $\beta_{v0,\text{ICV}}$ | $\mathcal{N}(0, 0.01)$ | |
| $\sigma_r$ | $\text{LogNormal}(a_0 + a_{\text{site}}, b)$ | such that $E[\sigma_r(a_0)] \approx \sigma_{r,\text{emp}}$, $Std[\sigma_r(a_0)] \approx \sigma_{r,\text{emp}}$ |
| $a_{\text{site}}$ | $\mathcal{N}(0, 0.5)$ | |
| $\beta(\text{site})$ | $\mathcal{N}(0, 0.3)$ | with $\sum_{site \in \text{sites}} \beta(site) = 0$ |

### 4 Standard Deviation of the Brain Structure Volumes

We plotted the standard deviation (SD) for age bins of 1 year throughout the adult lifespan. We observe that for all structures, except the ventricles, the SD is stable throughout the lifespan, with possibly a small tendency to shrink after the age of 60 years in the case of the Total gray matter volume and the supratentorial volume. If individuals have a large variation in trajectories, one would expect the SD to increase with time, not remain stable, as seen here. As discussed in the discussion, when the model is fitted to the data, this is likely some of the evidence that is taken into account when estimating the model parameters, and hence low variation in change is a likely scenario given the model.

### 5 Trace Plots

The trace plots of our MCMC runs, to show proper convergence and coverage of the posterior distribution (Geyer, 2011).

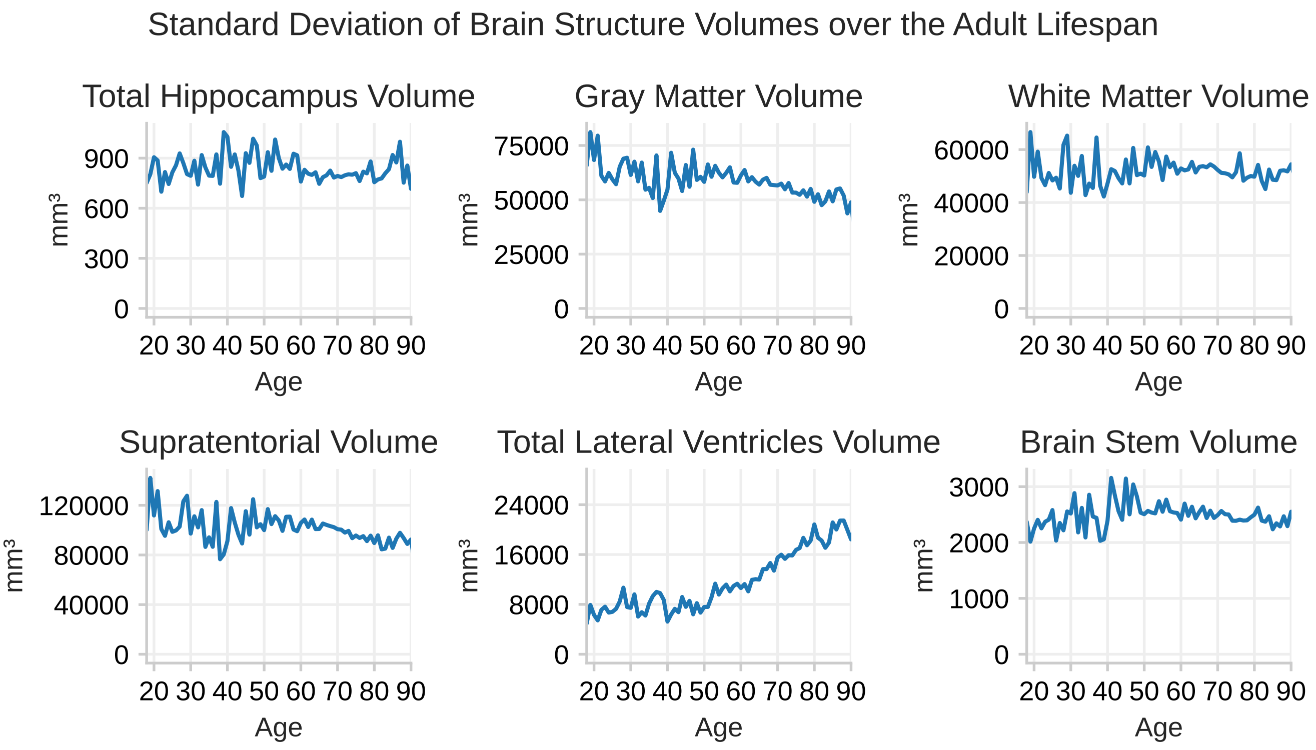


Figure 3: Standard deviation of the volume of the brain structures throughout the adult lifespan.

### 6 Empirical correlation

We computed the empirical correlation between volume and the estimated change rate. A delta estimate of the change rate will have a very high correlation with volume at any of the data points used to compute the delta estimate, which would mask the true correlation. We therefore computed an estimate of change for each session by using the time normalized difference from the session after to the session before. This ensures that the change estimate is independent of the volume measure. We then selected the first session for each subject with a valid change estimate and computed the correlation with volume. The result is shown in Fig. 5. The black lines in Fig. 5 of the main article show the correlation between the measured volume and the latent change rate. We expect the correlation in Fig. 5 to be of a higher magnitude while showing the same general trend, which is what we observe when comparing Fig. 5 to Appendix Fig. 5.

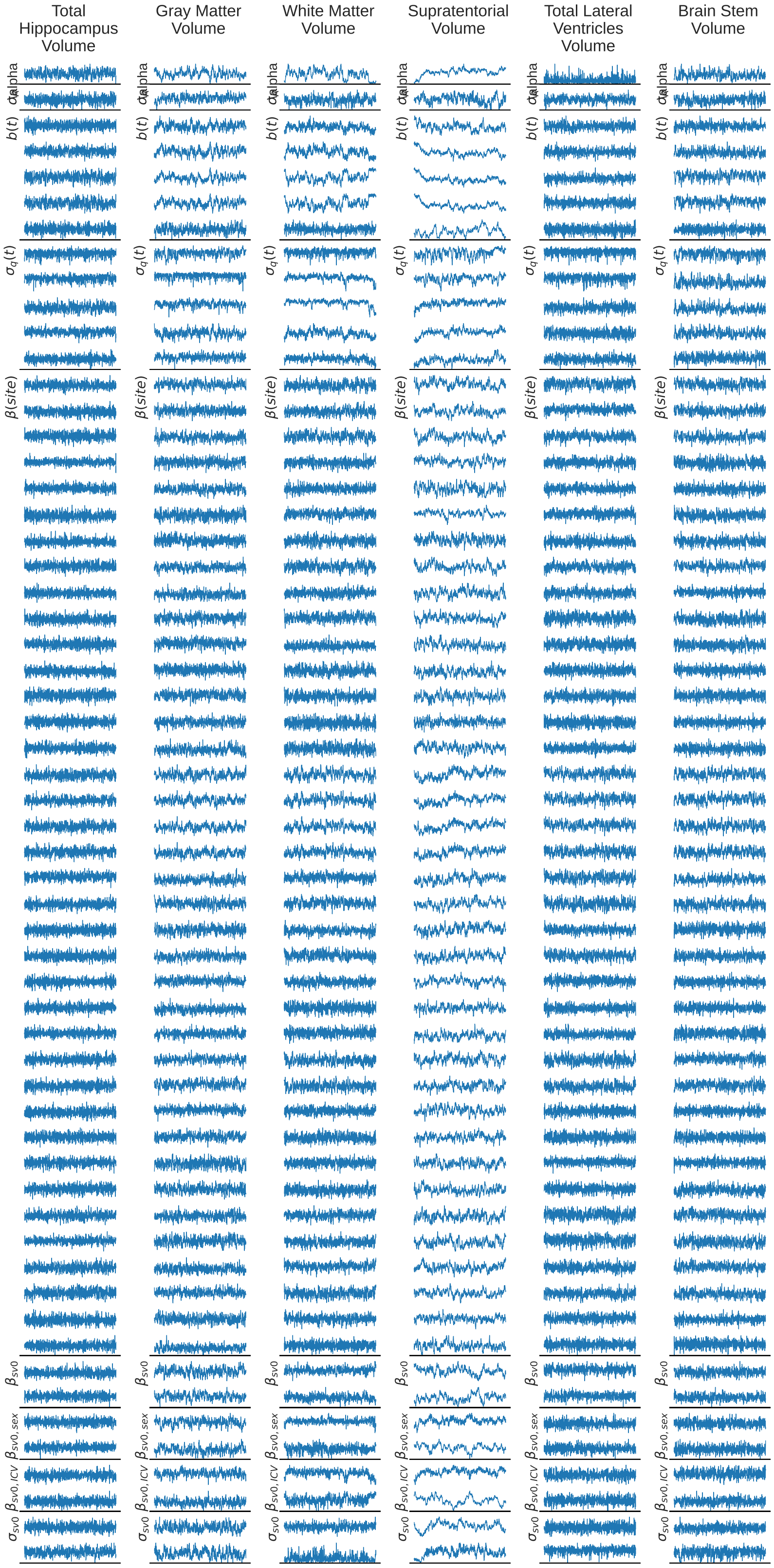


Figure 4: Trace plot of the MCMC runs. The parameters of the initial conditions are grouped together such that the first plot shows the latent structure volume parameter and the second the change rate.

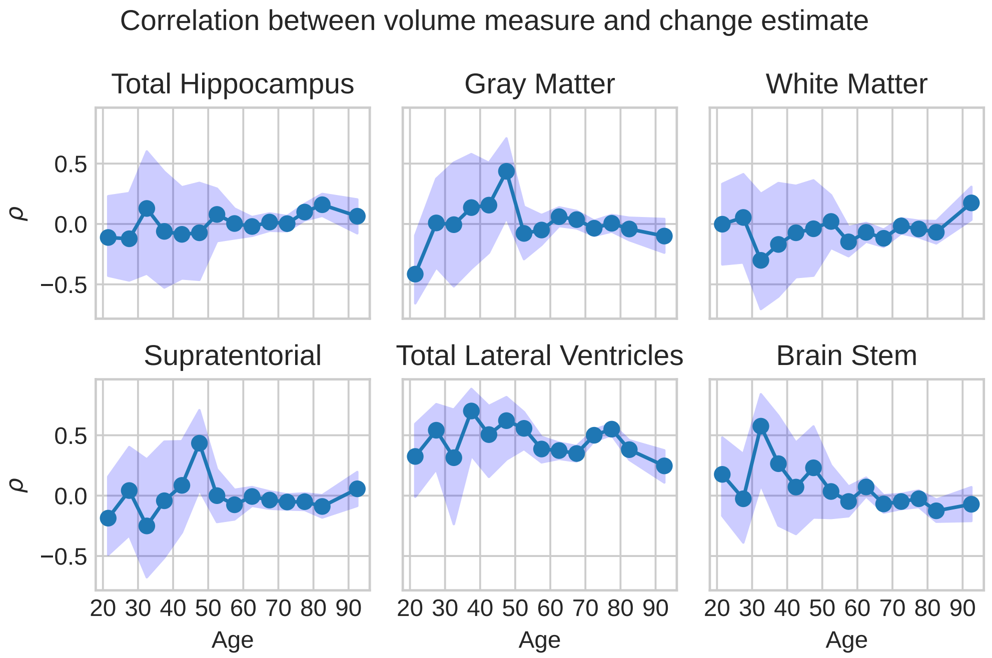


Figure 5: The correlation coefficient between the volume and measured change rate computed on the raw data in bins of 5 years. The shaded area show the 95 confidence interval of the correlation computed using the Fisher transformation.

### 7 Calibration Curves

In Fig. 6 the calibration curves of the fitted model are shown. These are created by plotting the cumulative probability distribution of the data points from the model on one axis while showing the proportions of data points that are below this threshold on the other. We find the model to be quite well calibrated. The ventricles is slightly further away from the optimal line, indicating that we are not fully able to capture the non-Gaussian probability density of this structure.

### 8 Bias Correction

Bias correction for each site is an integral part of the model. In Fig. 7 three quantities are shown. First, in blue, the z-scored volumes are shown, with the lines representing 95% of the data for each site. Next we have plotted in orange the z-scored volumes after the model has corrected for age, sex and ICV. Finally we show the the z-scored volumes after the bias correction is applied. For each structure we have calculated the mean absolute error (MAE) of the bias for each of these steps. We find that there are large differences in bias from each site and that our model successfully removes this bias.

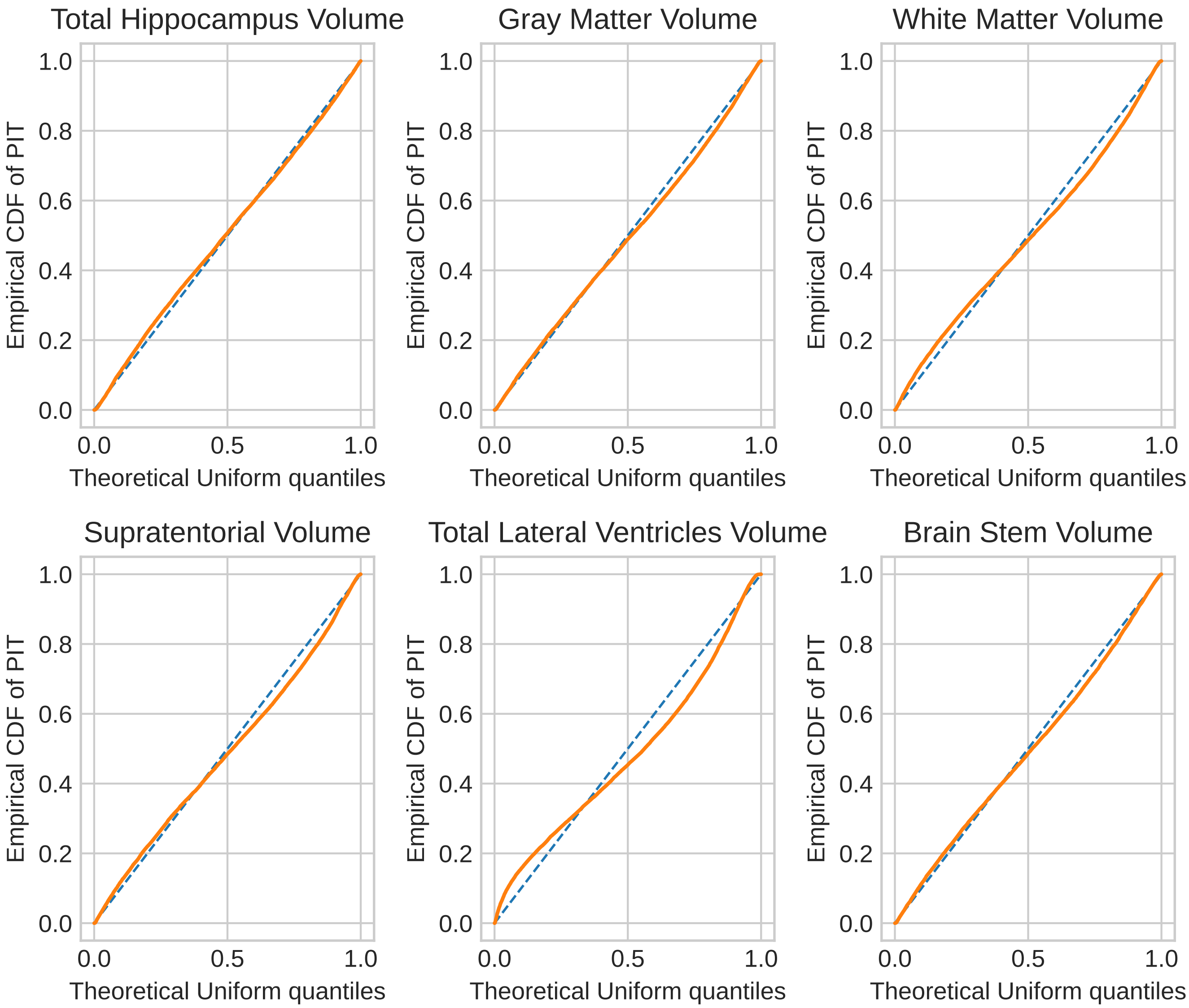


Figure 6: Calibration curve of the fitted model, using the mean of the posteriors.

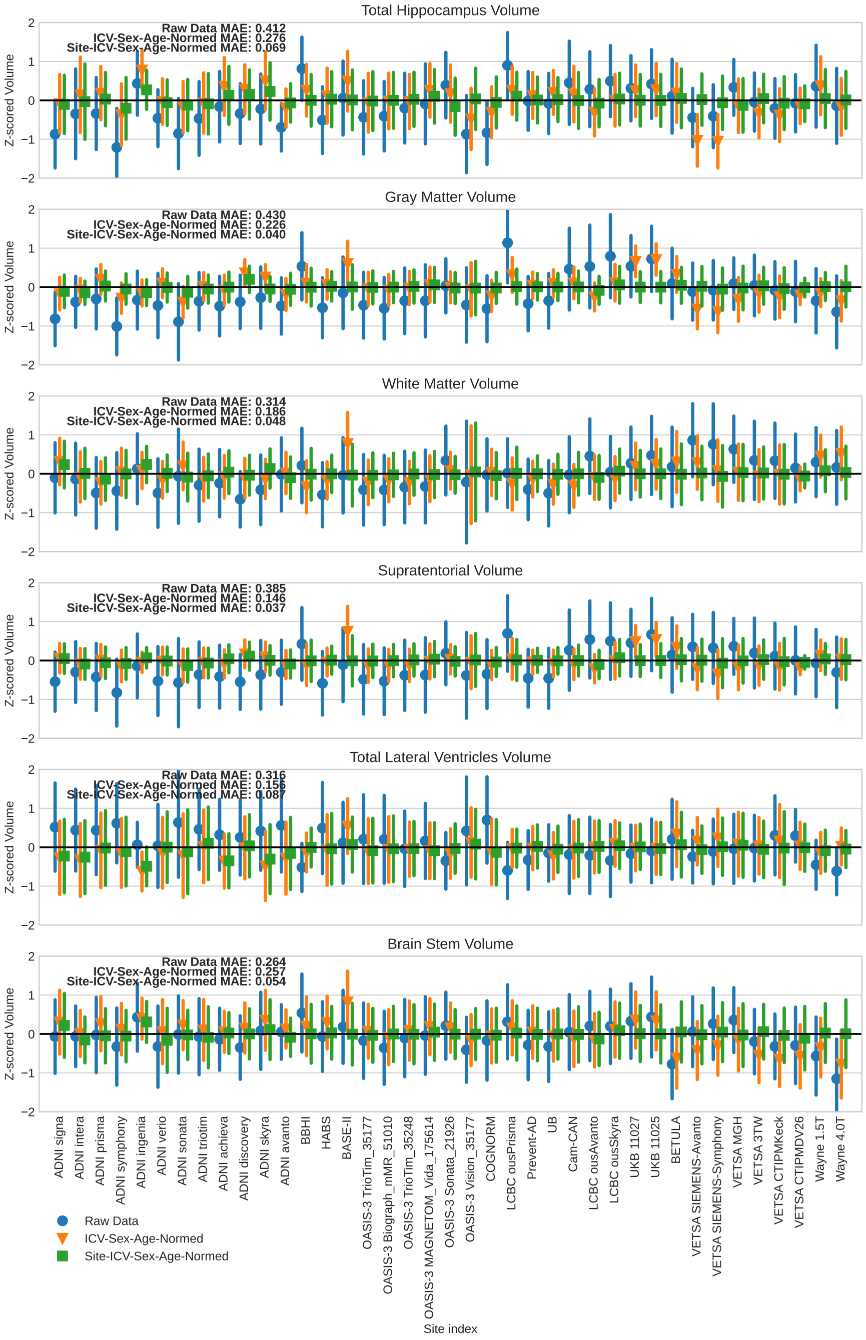


Figure 7: Results from bias correction.

### References

Burgmans, S., van Boxtel, M. P., Gronenschild, E., Vuurman, E., Hofman, P., Uylings, H. B., Jolles, J., & Raz, N. (2010). Multiple indicators of age-related differences in cerebral white matter and the modifying effects of hypertension. *NeuroImage*, *49*(3), 2083–2093.

Dagley, A., LaPoint, M., Huijbers, W., Hedden, T., McLaren, D. G., Chatwal, J. P., Papp, K. V., Amariglio, R. E., Blacker, D., Rentz, D. M., et al. (2017). Harvard aging brain study: Dataset and accessibility. *NeuroImage*, *144*, 255–258.

Geyer, C. J. (2011). Introduction to Markov chain Monte Carlo. *Handbook of Markov chain Monte Carlo*, *20116022*(45), 22.

Idland, A.-V., Sala-Llonch, R., Borza, T., Watne, L. O., Wyller, T. B., Brækhus, A., Zetterberg, H., Blennow, K., Walhovd, K. B., & Fjell, A. M. (2017). CSF neurofilament light levels predict hippocampal atrophy in cognitively healthy older adults. *Neurobiology of Aging*, *49*, 138–144.

Kennedy, K. M., & Raz, N. (2009). Aging white matter and cognition: Differential effects of regional variations in diffusion properties on memory, executive functions, and speed. *Neuropsychologia*, *47*(3), 916–927.

Kremen, W. S., Franz, C. E., & Lyons, M. J. (2019). Current status of the vietnam era twin study of aging (VETSA). *Twin Research and Human Genetics*, *22*(6), 783–787.

LaMontagne, P. J., Benzinger, T. L., Morris, J. C., Keefe, S., Hornbeck, R., Xiong, C., Grant, E., Hassenstab, J., Moulder, K., Vlassenko, A. G., et al. (2019). OASIS-3: Longitudinal neuroimaging, clinical, and cognitive dataset for normal aging and Alzheimer disease. *medRxiv*, 2019–12.

Miller, K. L., Alfaro-Almagro, F., Bangerter, N. K., Thomas, D. L., Yacoub, E., Xu, J., Bartsch, A. J., Jbabdi, S., Sotiropoulos, S. N., Andersson, J. L., et al. (2016). Multimodal population brain imaging in the UK Biobank prospective epidemiological study. *Nature Neuroscience*, *19*(11), 1523–1536.

Morris, J. C. (1993). The Clinical Dementia Rating (CDR) current version and scoring rules. *Neurology*, *43*(11), 2412–2412.

Mueller, S. G., Weiner, M. W., Thal, L. J., Petersen, R. C., Jack, C., Jagust, W., Trojanowski, J. Q., Toga, A. W., & Beckett, L. (2005). The Alzheimer's disease neuroimaging initiative. *Neuroimaging Clinics*, *15*(4), 869–877.

Nilsson, L.-G., Adolfsson, R., Bäckman, L., de Frias, C. M., Molander, B., & Nyberg, L. (2004). Betula: A prospective cohort study on memory, health and aging. *Aging Neuropsychology and Cognition*, *11*(2-3), 134–148.

Nyberg, L., Boraxbekk, C.-J., Sörman, D. E., Hansson, P., Herlitz, A., Kauppi, K., Ljungberg, J. K., Lövheim, H., Lundquist, A., Adolfsson, A. N., et al. (2020). Biological and environmental predictors of heterogeneity in neurocognitive ageing: Evidence from Betula and other longitudinal studies. *Ageing Research Reviews*, *64*, 101184.

Petersen, R. C., Aisen, P. S., Beckett, L. A., Donohue, M. C., Gamst, A. C., Harvey, D. J., Jack Jr, C., Jagust, W. J., Shaw, L. M., Toga, A. W., et al. (2010). Alzheimer's disease Neuroimaging Initiative (ADNI) clinical characterization. *Neurology*, *74*(3), 201–209.

Sajjad, M. U., Blennow, K., Knapskog, A. B., Idland, A.-V., Chaudhry, F. A., Wyller, T. B., Zetterberg, H., & Watne, L. O. (2020). Cerebrospinal fluid levels of interleukin-8 in delirium, dementia, and cognitively healthy patients. *Journal of Alzheimer's Disease*, *73*(4), 1363–1372.

Shafto, M. A., Tyler, L. K., Dixon, M., Taylor, J. R., Rowe, J. B., Cusack, R., Calder, A. J., Marslen-Wilson, W. D., Duncan, J., Dalgleish, T., et al. (2014). The Cambridge Centre for Ageing and Neuroscience (Cam-CAN) study protocol: A cross-sectional, lifespan, multidisciplinary examination of healthy cognitive ageing. *BMC neurology*, *14*, 1–25.

Tremblay-Mercier, J., Madjar, C., Das, S., Binette, A. P., Dyke, S. O., Étienne, P., Lafaille-Magnan, M.-E., Remz, J., Bellec, P., Collins, D. L., et al. (2021). Open science datasets from PREVENT-AD, a longitudinal cohort of pre-symptomatic Alzheimer's disease. *NeuroImage: Clinical*, *31*, 102733.

Walhovd, K. B., Fjell, A. M., Westerhausen, R., Nyberg, L., Ebmeier, K. P., Lindenberger, U., Bartrés-Faz, D., Baaré, W. F., Siebner, H. R., Henson, R., et al. (2018). Healthy minds 0–100 years: Optimising the use of European brain imaging cohorts ("Lifebrain"). *European Psychiatry*, *50*, 47–56.

Walhovd, K. B., Krogsrud, S. K., Amlien, I. K., Bartsch, H., Bjørnerud, A., Due-Tønnessen, P., Grydeland, H., Hagler Jr, D. J., Håberg, A. K., Kremen, W. S., et al. (2016). Neurodevelopmental origins of lifespan changes in brain and cognition. *Proceedings of the National Academy of Sciences*, *113*(33), 9357–9362.
